# Single-molecule mitochondrial DNA imaging reveals heteroplasmy dynamics shaped by developmental bottlenecks and selection in different organs *in vivo*

**DOI:** 10.1101/2025.01.24.634671

**Authors:** Rajini Chandrasegaram, Antony M. Hynes-Allen, Beitong Gao, Abhilesh Dhawanjewar, Michele Frison, Stavroula Petridi, Patrick F. Chinnery, Hansong Ma, Jelle van den Ameele

## Abstract

Mitochondrial DNA (mtDNA) occurs in many copies per cell, with cell-to-cell variability in mutation load, known as heteroplasmy. Developmental and age-related expansion of pathogenic mtDNA mutations contributes to mitochondrial and neurodegenerative disease pathogenesis. Here, we describe an approach for *in situ* sequence-specific detection of single mtDNA molecules (mtDNA-smFISH). We apply this method to visualize and measure *in situ* mtDNA and heteroplasmy levels at single-cell resolution in whole-mount *Drosophila* tissue and cultured human cells. In *Drosophila*, we identify a somatic mtDNA bottleneck during neurogenesis. This amplifies heteroplasmy variability between neurons, as predicted from a mathematical bottleneck model, predisposing individual neurons to a high mutation load and degeneration. However, both during neurogenesis and oogenesis, mtDNA segregation is accompanied by purifying selection, promoting wild-type over mutant mtDNA. mtDNA-smFISH thus elucidates novel mechanisms whereby developmental cell-fate transitions, accompanied by changes in cell morphology, behaviour and metabolism, will shape disease-relevant and tissue-specific transmission and selection of mtDNA mutations.

## Introduction

Unlike the nuclear genome, which is haploid or diploid in most animal cells, the mitochondrial genome (mtDNA) occurs in up to many thousands of copies per cell. Most human cells carry a mixture of mutant and wild-type (WT) mtDNA (heteroplasmy) ^1–3^, with considerable cell-to-cell variability in mutation load ^4,5^. During development and ageing, mtDNA mutations can clonally expand to high levels in single cells, causing defects in energy production and metabolism, and eventually cell death ^6–8^. Expansion of pathogenic mtDNA mutations follows striking tissue- and cell-type-specific patterns during development and aging ^7,9^, and is the main cause of inherited and sporadic adult-onset mitochondrial diseases ^10^. Clonal amplification of mtDNA mutations has also been implicated in common neurodegenerative disorders like Alzheimer’s and Parkinson’s disease ^11,12^, or in cancer, where certain somatic mtDNA variants reach homoplasmy due to positive selection ^13^. In *Drosophila*, deleterious mtDNA mutations have also been shown to increase within individuals over time ^14–16^, contributing to developmental and aging phenotypes, and even lethality ^16–22^.

Variability in heteroplasmy levels between cells within a tissue, or between a mother and her off-spring, can be caused by vegetative segregation, further accelerated by random drift in heteroplasmy levels, due to the ongoing destruction and replication of mtDNA, independent of the cell cycle (relaxed replication) ^4,6^. These processes may be modulated by purifying selection or selfish propagation, resulting respectively in a decrease or an increase of mutant mtDNA heteroplasmy levels ^8,15,23^. A special example occurs between mothers and their offspring, where heteroplasmy shifts are further accentuated by a genetic mtDNA bottleneck, due to drastic mtDNA copy number (CN) reduction in single cells during maternal germline development ^24–29^. Although well studied in the germline, little is known about segregation of heteroplasmy in somatic tissues during development and aging. Nonetheless, this is likely to play a crucial role in determining mutational burden during life. For example, many developmental and homeostatic cell-fate transitions are accompanied by profound changes in cell volume, morphology, and metabolism, but it remains unknown how these affect the mtDNA, and may amplify shifts in heteroplasmy levels.

Current explanations of heteroplasmy shifts are mainly based on modelling of data from whole-tissue analysis ^30^ or single-cell sequencing and genotyping of isolated cells outside their *in vivo* context ^4,5,31–34^. A major limitation of these approaches is that they mostly rely on tissue dissociation, losing spatial information and limiting analysis of subcellular mechanisms.

Here, we develop and validate a novel approach for *in situ* visualization of heteroplasmic mtDNA variants (mtDNA-smFISH) based on single-molecule *in situ* hybridisation (smFISH) combined with enzyme-free hybridisation chain reaction (HCR) for signal amplification. We apply this method to detect mtDNA heteroplasmy *in situ* in whole-mount tissue of an established *Drosophila* model of stable heteroplasmy, and in cultured human cybrid cells carrying a heteroplasmic single large-scale mtDNA deletion. In *Drosophila*, mtDNA-smFISH confirmed the presence of purifying selection against mutant mtDNA during oocyte development. In addition, we provide evidence for the existence of a somatic bottleneck during neurogenesis in the developing larval brain. This amplifies cell-to-cell variability in heteroplasmy levels between different progeny derived from the same neural stem cell (NSC), thus predisposing subsets of neurons and glia to high levels of deleterious mtDNA mutations, leading to mitochondrial dysfunction and disease.

### Design

Current mtDNA imaging methods either lack the resolution to distinguish single mtDNA molecules ^35,36^, have low sensitivity ^14^, depend on mtDNA replication ^37^, have not been used on whole-mount tissues ^38–40^, or are not compatible with the genetics of multicellular organisms ^41^. Development of *in vivo* models and tools to measure, track and modulate heteroplasmy in somatic tissues is required to understand the cellular and molecular processes that cause somatic heteroplasmy variation and tissue-specific phenotypes of mitochondrial disease, and answer long-standing questions in the field of mitochondrial/mtDNA biology ^42^.

In order to develop an approach for *in situ* detection of heteroplasmic mtDNA variation in whole-mount tissues, we took advantage of recent advances in single-molecule Fluorescent *In situ* Hybridisation (smFISH) combined with Hybridisation Chain Reaction (HCR). smFISH-HCR is a well-established technique for the detection of mRNA in whole-mount tissues ^43–46^. It combines short single-stranded DNA probes complementary to target nucleotide sequences with enzyme-free signal amplification based on fluorescently labelled metastable DNA hairpins (**Figure 1A)**. Third-generation smFISH-HCR achieves high signal-to-background ratios by relying on pairs of short (25bp) single-stranded DNA probes that together recognize a 52bp mRNA target sequence, and initiate HCR only when bound in close proximity and correct orientation ^43^ (**Figure 1B**). This allows *in situ* single-molecule detection with as few as 5 probe pairs or less ^46^. In order to detect mtDNA instead of mRNA, we established a protocol of dsDNA denaturation in whole-mount tissue through heating of formaldehyde- and methanol-fixed tissue to 95°C in 70% formamide (see methods). This allows double-stranded mtDNA to become single-stranded, so the probes can anneal to their complementary mtDNA-sequence *in situ*.

**Figure 1:**
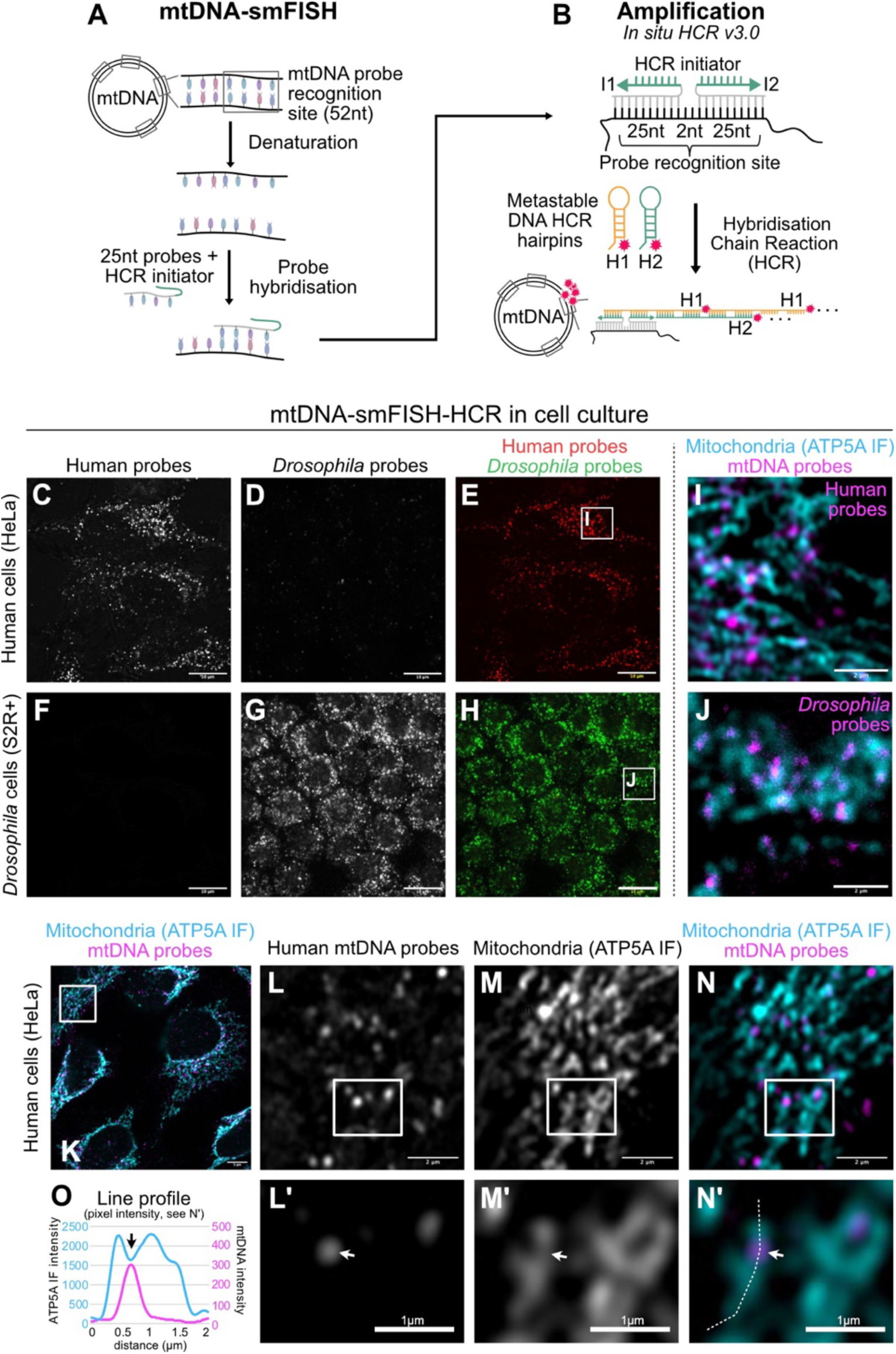
Single-molecule Fluorescent *In situ* hybridisation of mtDNA (mtDNA-smFISH) in cell culture. (A) Pairs of 25-nucleotide (nt) probes were designed against 52nt mtDNA target sequences, and used for in situ hybridisation after denaturation. (B) Signal amplification relied on metastable DNA hairpins recognising an initiator sequence formed when a probe pair hybridises on the target sequence, initiating an enzyme-free hybridisation chain reaction (HCR). (C-H) mtDNA-smFISH in human HeLa (C-E) or *Drosophila* S2R+ (F-H) cell culture with probes against human (C,F,E,H) or *Drosophila* (D,G,E,H) mtDNA. (I,J) zoomed areas boxed in (E,H) showing mtDNA-smFISH (magenta) followed by co-immunostaining (cyan) against the mitochondrial marker ATP5A. (K-N) Airyscan high-resolution imaging of HeLa cells after mtDNA-smFISH with human mtDNA probes (magenta) with co-immunostaining of ATP5A (cyan), with higher magnification of boxed regions (L-N) in (L’-N’). (O) Line profile analysis of signal along the hatched line in N’. Scale bars are 10μm (C-H), 5μm (K), 2μm (I,J,L-N) or 1μm (L’-N’). See also Figure S1; Table S1.

## Results

### Single-molecule Fluorescent *In Situ* Hybridisation of mtDNA (mtDNA-smFISH)

We first designed sets of 12 and 10 probe pairs to specifically detect human or *Drosophila* mtDNA respectively (**Figure S1A,B**; **Table S1**). We used these to conduct mtDNA-smFISH on human or *Drosophila melanogaster* cells cultured *in vitro*. Each probe set generated a distinct cytoplasmic punctate staining pattern in the cells of its corresponding species (human or *Drosophila*) (**Figure 1C-H**). In contrast, when the same probe sets were applied to the converse cell types (i.e. human probes on *Drosophila* cells and vice versa) no signal was detected (**Figure 1D,F**), indicating specificity of the probes for the respective mtDNA molecules.

To further confirm that the punctate pattern did correspond to mtDNA, we conducted fluorescent immunohistochemistry (IHC) with antibodies against mitochondrial epitopes that withstand the harsh denaturation conditions of mtDNA-smFISH. Both in human HeLa and *Drosophila* S2R+ cells, co-IHC of ATP-synthase, situated at the inner mitochondrial membrane, together with mtDNA-smFISH showed a clear overlay of smFISH puncta with the mitochondrial network (**Figure 1I-N**). Notably, we often observed a gap in the ATP-synthase staining observed at the site of mtDNA-smFISH (**Figure 1L-O**). This may be due to steric hindrance from the HCR hairpins, preventing antibody-access ^47,48^. However, a similar lack of overlap between ATP-synthase IHC and mtDNA nucleoids has been described previously using other approaches ^49–51^, suggesting that mtDNA nucleoids may be situated away from mitochondrial cristae or the ATP-synthase complex within those cristae.

### mtDNA-smFISH in whole-mount tissue

We next applied mtDNA-smFISH to whole-mount *Drosophila* larval brains and observed a similar cytoplasmic punctate staining pattern throughout the entire brain (**Figure 2A,B**). No signal was observed in conditions without probes or amplifiers (**Figure S2A-F**), and probes recognized mtDNA from either *D. yakuba* (Dyak) or *D. melanogaster* (Dmel) strains, as they were designed against conserved regions of the two mitochondrial genotypes (**Figure S2G-I**). To estimate sensitivity, we quantified smFISH puncta in single NSCs, which can be identified within the larval ventral nerve cord (VNC) by their large nuclear size and expression of GFP under control of a NSC-specific GAL4 ^52,53^, and compared this to standard mtDNA copy number (CN) quantification by single-cell digital droplet PCR (ddPCR) of FACS-sorted nuclear GFP-labelled NSCs (**Figure S3;4**). The average NSC mtDNA CN detected by smFISH was 268.5±37.1 (mean±sd) (**Figure 2C**), and 292.5±140.4 (mean±sd) by ddPCR (**Figure 2D**), indicating >90% efficiency of mtDNA-smFISH to detect mitochondrial nucleoids. The difference between both techniques likely reflects mtDNA replication occurring in some nucleoids, which are therefore expected to have >1 copy of mtDNA per mtDNA-smFISH punctum ^54^. Importantly, these quantifications indicate that also *in vivo*, and in non-mammalian species, the majority of nucleoids harbor only a single copy of mtDNA, which bears striking resemblance to previous observations from *in vitro* studies with cultured mammalian cells ^54,55^.

**Figure 2:**
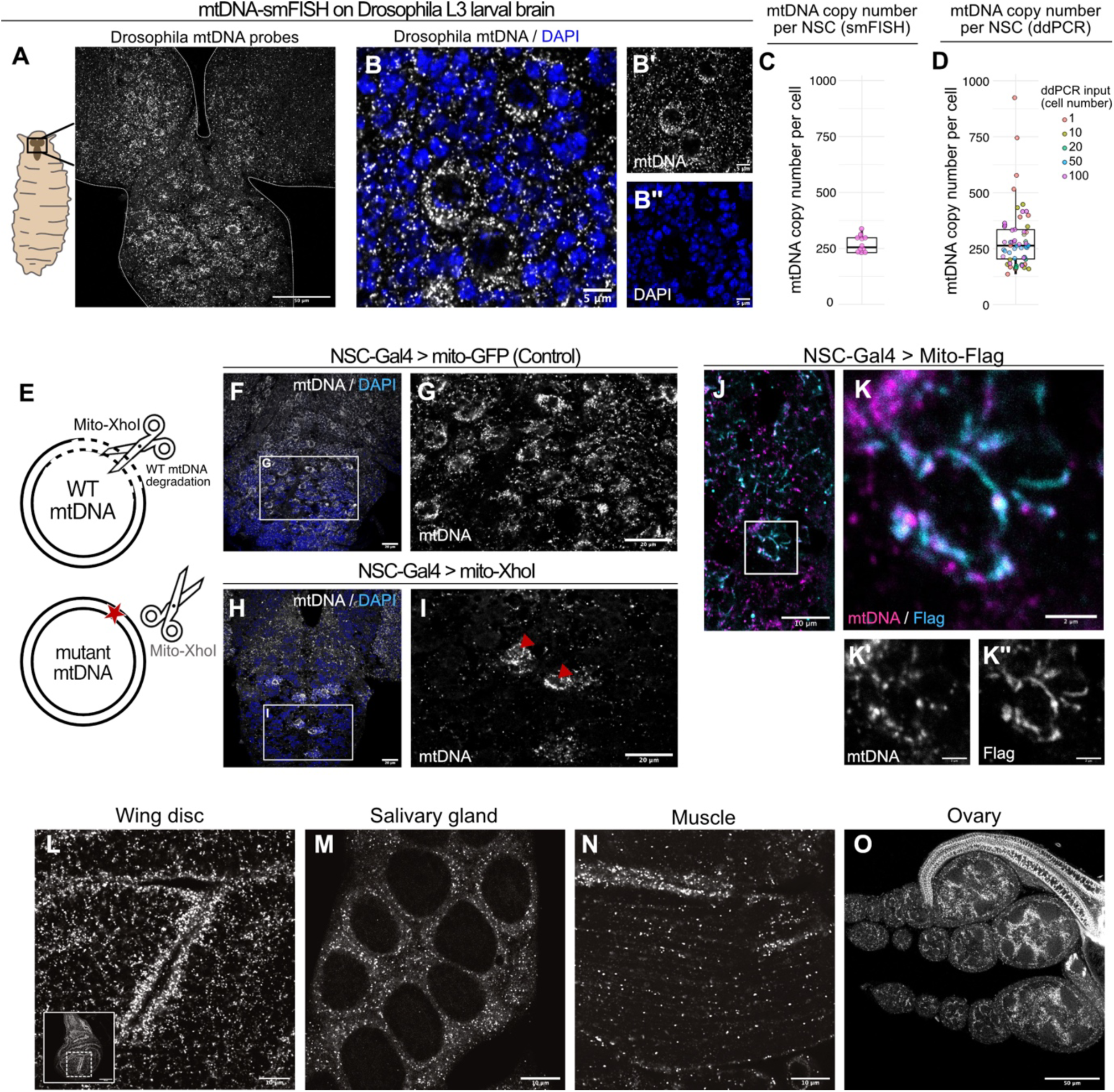
mtDNA-smFISH in whole-mount *Drosophila* tissue. (A,B) mtDNA-smFISH (gray) in *Drosophila* third instar larva brain and co-labelling with DAPI (blue) to mark nuclei. Single channels of B are in B’, B’’. (C,D) mtDNA quantification in single NSCs by mtDNA-smFISH (C) or ddPCR (D), with each dot representing mtDNA CN in a single NSC. (E) Schematic of mito-XhoI activity resulting in digestion and degradation of wild-type (WT) but not mutant mtDNA. (F-I) Mito-GFP control (F,G)t or mito-XhoI (H,I) expression in NSCs (Worniu-GAL4) of homoplasmic WT *D. melanogaster* larval brains, followed by mtDNA-smFISH (gray) and DAPI counterstaining (blue). Red arrowheads indicate NSCs that do not express Worniu-GAL4 and therefore retain WT mtDNA. (J,K) mtDNA-smFISH (magenta) and immunostaining (cyan) against Flag in NSCs expressing a mitochondrial-targeted Flag-tagged transgene. Single channels of K are in K’, K’’. (L-O) mtDNA-smFISH across various *Drosophila* tissues. Scale bars are 50μm (A,L,O), 20μm (F-I), 10μm (J,M,N), 5μm (B) or 2μm (K). See also Figure S2,3,4.

Expression of a mitochondrial-targeted restriction enzyme, mito-XhoI in *Drosophila* cells was previously shown to linearise mtDNA through digestion at a single restriction site (**Figure 2E**), resulting in mtDNA degradation and depletion ^56^. When we expressed mito-XhoI specifically in NSCs and their progeny with *Worniu*-GAL4, the mtDNA-smFISH signal was lost, apart from in cells not expressing mito-XhoI (**Figure 2F-I**). Co-IHC of a mitochondrial-targeted Flag-tagged transgene expressed in NSCs showed clear overlap with mtDNA-smFISH signals (**Figure 2J,K**), further confirming the specificity of mtDNA-smFISH. Moreover, mtDNA-smFISH performed reliably in other developing and adult *Drosophila* tissues, including the wing imaginal discs, larval salivary gland, thoracic muscle and ovaries (**Figure 2L-O**).

### Immunofluorescence combined with mtDNA-smFISH

Although we could successfully conduct anti-ATP-synthase (**Figure 1I-N**) and anti-Flag-tag (**Figure 2J-K**) IHC after the harsh denaturation conditions of mtDNA-smFISH, this was not the case for most other epitopes, such as GFP (**Figure S5**), did not tolerate this treatment as they change conformation upon fixation, heating or formamide treatment. Although specific anti-heat denatured antibodies exist for GFP ^57^, allowing anti-GFP immunostaining in whole-mount tissues after denaturation (**Figure 3A-D**), such reagents are not available for most proteins or epitopes. We therefore developed a protocol that includes immunostaining with biotin-labelled antibodies prior to denaturation (**Figure 3E**), as biotin remains intact during the denaturation step. When combined with a second round of fixation after primary antibody incubation (see methods), this allows visualization of the primary antibody binding pattern with a streptavidin-conjugated fluorophore post-mtDNA-smFISH (**Figure 3F,G**).

**Figure 3:**
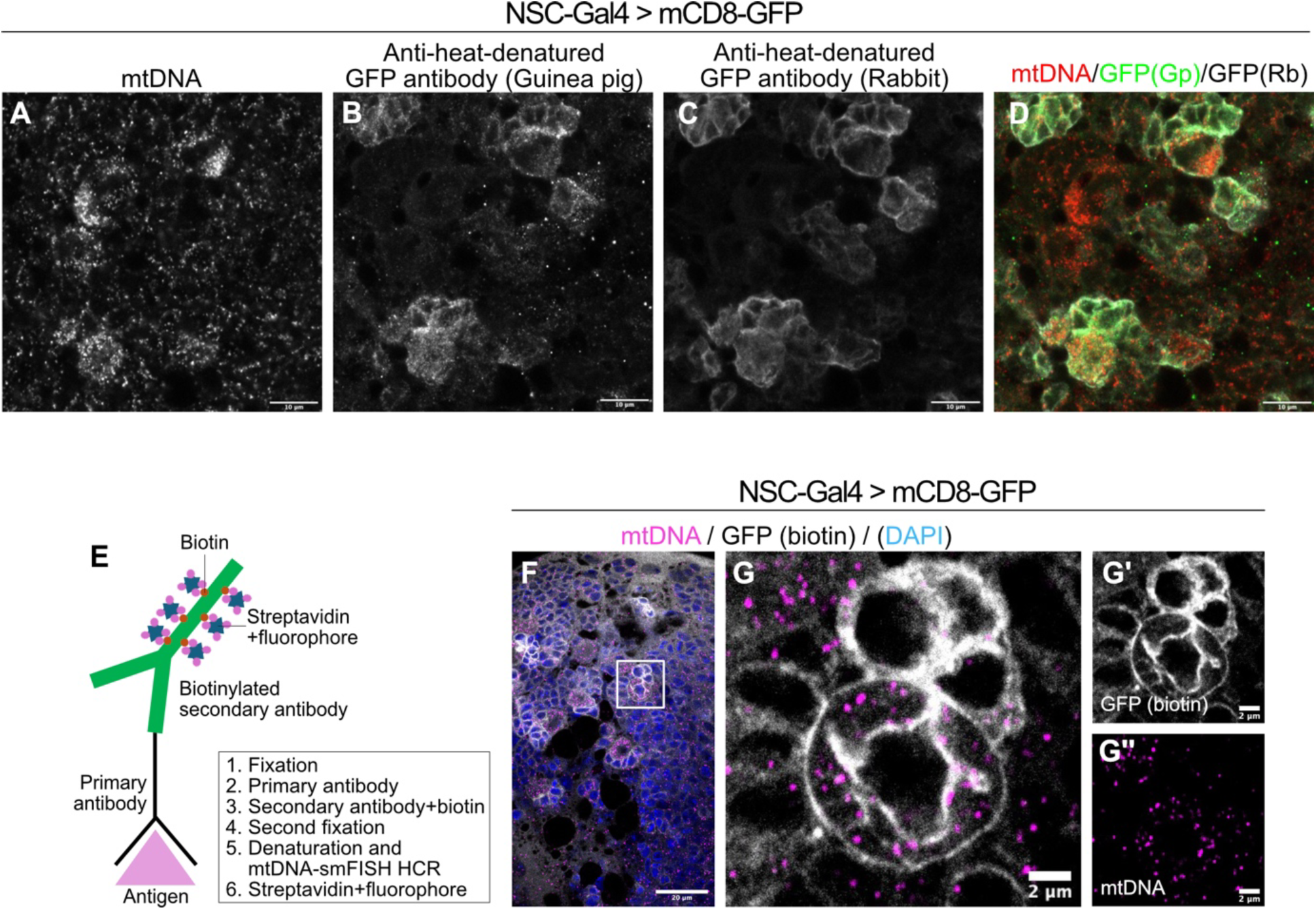
Immunofluorescence combined with mtDNA-smFISH. (A-D) mtDNA-smFISH (A) with immunostaining using anti-heat denatured GFP antibodies raised in rabbit (B), or guinea pig (C) to mark NSCs (Worniu-GAL4 > UAS-mCD8-GFP) in the *Drosophila* larval brain. (E) Schematic of biotin-labelled antibody immunostaining to circumvent denaturation of antigens. (F,G) mtDNA-smFISH (magenta) combined with biotin-labelled antibody staining of mCD8-GFP in NSCs (Worniu-GAL4; gray) and DAPI counterstain (blue). Scale bars are 20μm (F), 10μm (A-D), or 2μm (G-G’’). See also Figure S5.

### mtDNA-smFISH to detect *in situ* mtDNA heteroplasmy *in vivo*

Having established mtDNA-smFISH for sequence-specific *in situ* detection of single mtDNA molecules, we next asked whether this approach would allow us to distinguish different mtDNA molecules within the same cell. mtDNA deletions have previously been detected *in situ* in cultured human cells by FISH using long probes ^36^. We adapted mtDNA-smFISH for the same purpose, to detect mtDNA heteroplasmy in human cybrid cells carrying a mixture of wild-type (WT) mtDNA and mtDNA molecules with a single large deletion ^58^. Specifically, we designed one set of 12 probe pairs to specifically label full-length mtDNA molecules by targeting the mtDNA deletion sequence, and compared this to a probe set recognizing all (i.e. deleted and full-length) mtDNA molecules (**Figure 4A; Figure S1D,G,H; Table S1**). Using both human probe sets recognising either the deleted or non-deleted sections of the human mtDNA for mtDNA-smFISH, we observed pronounced cell-to-cell variation in the amount of full-length mtDNA molecules (**Figure 4B-D**), reminiscent of single-cell sequencing findings in blood cells derived from patients with single mtDNA deletions ^31^.

**Figure 4:**
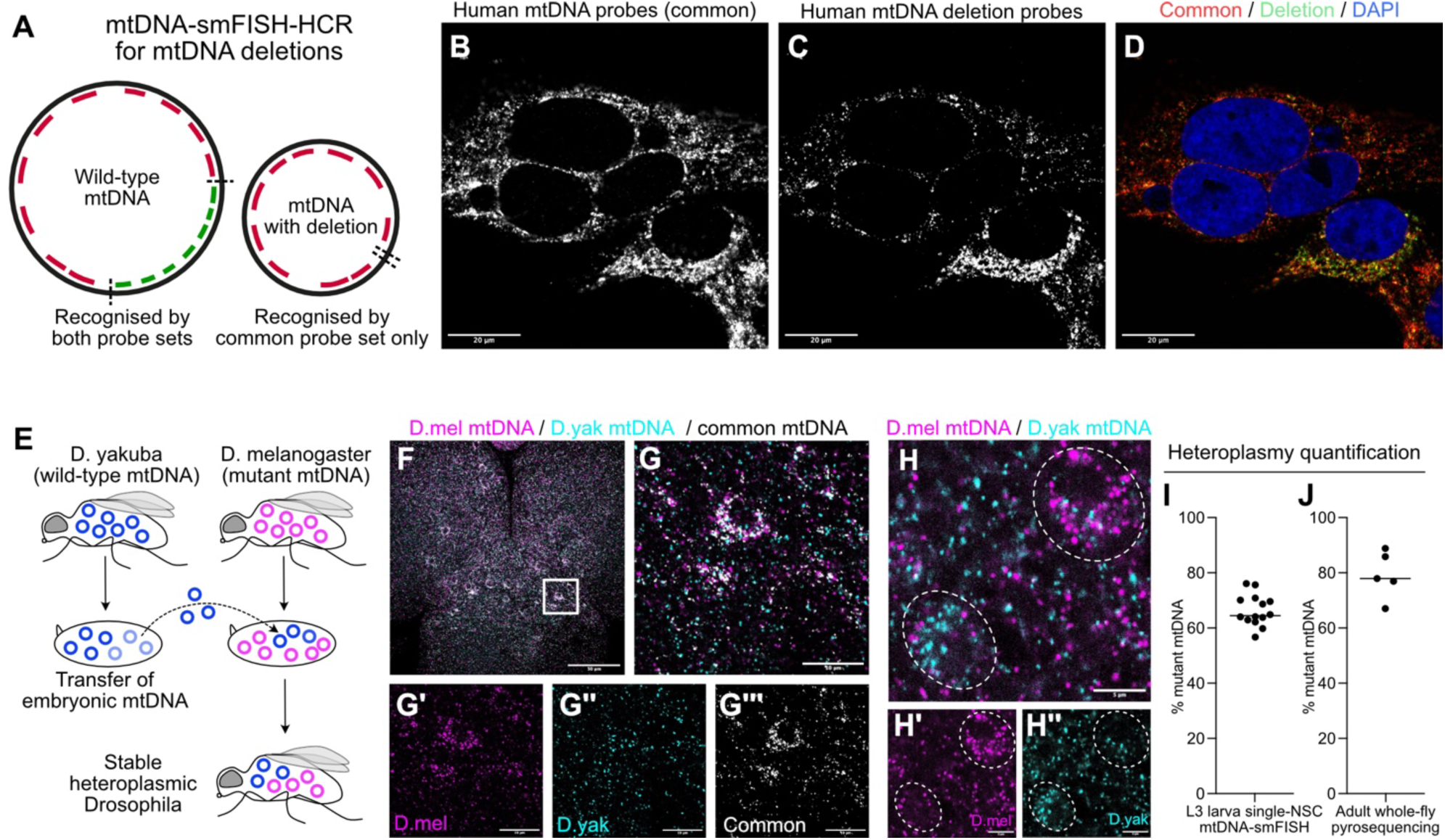
Visualizing heteroplasmy with mtDNA-smFISH. (A) Schematic of mtDNA-smFISH for detection of single mtDNA deletions. (B-D) mtDNA-smFISH in cybrid cells in culture with probes against all human mtDNA molecules (B, red) or only against the region deleted in the converse mtDNA molecule (C, green), with DAPI counterstain (blue). Note that mtDNA deletion probes only recognize the molecules that do not carry the deletion. (E) Generating heteroplasmic Drosophila through mtDNA-transfer between different species. (F-H) mtDNA-smFISH using probes targeting only *D. melanogaster* (mutant, magenta, G’,H’), only *D.yakuba* (WT, magenta, G’’,H’’) or mtDNA from both species (gray, G’’’) in heteroplasmic *Drosophila* brains. Dashed outlines indicate NSCs. (I,J) Heteroplasmy levels (% mutant mtDNA) in single NSCs of L3 larval brains quantified after mtDNA-smFISH (I) or in whole adult flies determined by pyrosequencing (J). Scale bars are 50μm (F), 20μm (B-D), 10μm (G) or 5μm (H). See also Figure S6,7,8; Table S1.

To study mtDNA heteroplasmy *in vivo*, we took advantage of a heteroplasmic *D. melanogaster* (Dmel) fly line that stably transmits a temperature-sensitive lethal mtDNA mutation in the CoI subunit of the cytochrome oxidase complex (*mt:CoI^T300I^*) together with a 9bp deletion (*mt:ND2^del^*^1^) in the ND2 subunit of Complex I ^56^. The *mt:CoI^T300I^* mutation impairs cytochrome oxidase activity at higher temperature (29°C), causing lethality at late pupal stages in homoplasmic flies ^21,59^. Introducing WT mtDNA from *D. yakuba* (Dyak) species through germ-plasm transfer ^60^ rescues viability in *mt:CoI^T300I^* flies ^15,35,61^. As a result, these *D. melanogaster* flies (i.e. with a *D. melanogaster* nuclear genome) carry both the Dmel *mt:COI^T300I^* mutant and Dyak WT mtDNA at high levels of heteroplasmy (**Figure 4E**). We designed mtDNA-smFISH probes to distinguish Dmel (carrying the *mt:CoI^T300I^* mutation) from Dyak (WT) mtDNA (**Figure S1C,E; Table S1**). The main sequence divergence between Dmel and Dyak mtDNA is in the non-coding D-loop, allowing FISH with long probes ^35^, but precluding single-molecule analysis. Due to its repetitive and AT-rich nature, this non-coding region is not amenable to smFISH with small probes. We therefore designed 7 (Dmel) and 5 (Dyak) probe-pairs across the most divergent coding regions characterised by small indels or multiple single-nucleotide polymorphisms (**Figure S1C**). mtDNA-smFISH in heteroplasmic larval brains yielded clear punctate cytoplasmic staining for both probe sets (**Figure 4F,G**), albeit weaker than mtDNA-smFISH with the common probe set consisting of 10 probe pairs (**Figure S2;S6**).

As with the other probe sets, no signal was detected in conditions without probes or amplifiers (**Figure S6A-L**). Testing specificity on brains from WT homoplasmic Dmel or Dyak flies (i.e. only containing WT *D. melanogaster* or WT *D. yakuba* mtDNA), showed that the Dmel probe set produced no signal in Dyak tissue (**Figure S6M-T**). Some punctate staining was observed with the Dyak probe set on homoplasmic Dmel tissues (**Figure S6O,S**), which was not due to a specific probe pair from the Dyak probe set (**Figure S7**). This background was considerably weaker than the signal obtained in homoplasmic Dyak tissue (**Figure S6O,S,R,T**), allowing background correction during subsequent image analysis.

We quantified heteroplasmy levels in single NSCs by mtDNA-smFISH, and observed notable cell-to-cell variability in heteroplasmy levels, even between NSCs of the same brain (**Figure 4H,I**). On average, NSC heteroplasmy levels measured by mtDNA-smFISH were slightly lower than the levels obtained by pyrosequencing of whole adult flies of the same genotype (**Figure S8C-G; Figure 4J**), consistent with previously described age-dependent increase of mutant mtDNA ^16^. Large NSC-to-NSC heteroplasmy variability was also observed using pyrosequencing of mtDNA from FACS-sorted single NSCs with a different nuclear genomic background (Dpn::GFP; **Figure S8E**). Because mtDNA-smFISH allows direct *in situ* visualization of heteroplasmy, it opens the possibility to investigate heteroplasmy shifts across different cells and tissues, and eventually uncovering the underlying cellular and molecular processes.

### Purifying selection in the *Drosophila* female germline

Previous studies indicate that mutant mtDNA molecules are subject to purifying selection during oogenesis in *Drosophila* ^35,62–65^. These conclusions were mostly based on qPCR-based analysis of mtDNA in homogenised tissue from ovaries or offspring from heteroplasmic female *Drosophila* ^35,62–64^, or on whole-tissue analysis with long FISH probes ^35^. However, single-cell, single-molecule analysis of this process has not been possible so far. We used mtDNA-smFISH to label Dmel or Dyak mtDNA in individual cells and compartments of heteroplasmic *Drosophila* ovaries (**Figure 5A,B**), and conducted automated 3D quantification of mtDNA with FISH-Quant ^66^. This showed a small but significant decrease in the percentage of mutant Dmel mtDNA between the regions of the germarium corresponding to stages 1+2a (53.1±3.9%, mean±sd) or 2b (52.0±6.0%, mean±sd) and stage 3 (48.5±2.7%, mean±sd) (p=0.02 and p=0.03 respectively) (**Figure 5C-D**). Interestingly, when imaging entire ovaries, we also noticed instances of large cell-to-cell variability in heteroplasmy levels in the nurse cell compartments (**Figure 5E**). This likely indicates that the mother-to-offspring bottleneck caused by low mtDNA levels in germline stem cells (GSCs) ^29^ and subsequent rapid replication during oogenesis and nurse cell formation, also enhances cell-to-cell variability in heteroplasmy levels for non-oocyte GSC-derived cell-types.

**Figure 5:**
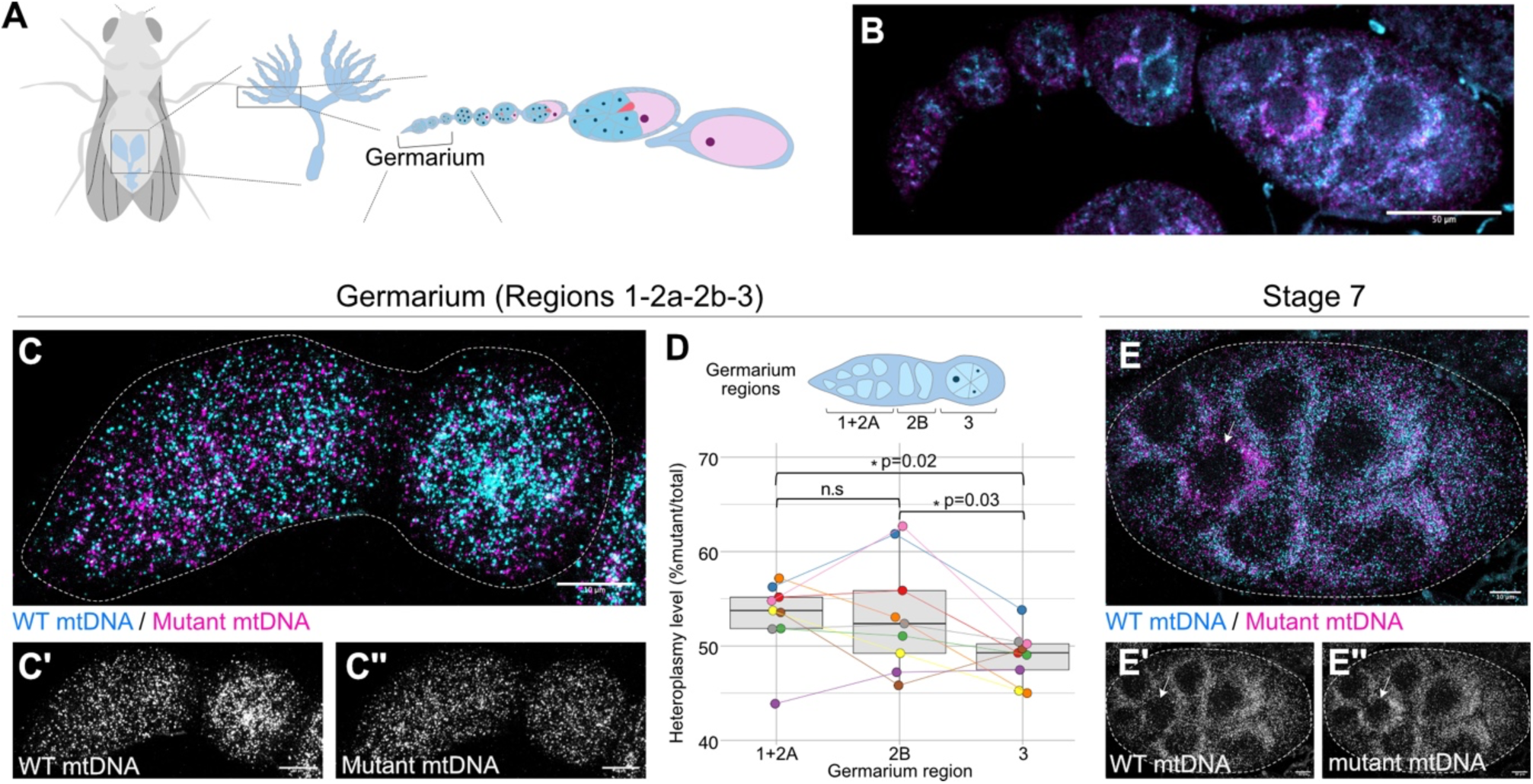
Purifying selection in the *Drosophila* female germline. (A,B) Schematic of *Drosophila* oocyte development, with developing oocytes arranged in linear arrays consisting of 15-20 ovarioles each. (B,C) Representative images of mtDNA-smFISH using probes targeting only *D. melanogaster* (mutant, magenta, C’’) or *D.yakuba* (WT, cyan, C’) mtDNA across the germline (B) or the germarium only (C, outline). (D) The germarium is divided in three regions that define stages of germ cell development. Quantification of mtDNA-smFISH puncta in regions 1+2a, 2b and 3, with dots indicating single germaria (paired t-test). (E) Representative image of mtDNA-smFISH using probes targeting only *D. melanogaster* (mutant, magenta, E’’) or *D.yakuba* (WT, cyan, E’) mtDNA in stage 8 *Drosophila* egg chamber nurse cells. Arrow indicates a nurse cell with high heteroplasmy level. Scale bars are 50μm (B), 10μm (C,E).

### A somatic bottleneck during *Drosophila* neurogenesis

During development, many instances occur in which large stem cells undergo asymmetric division to self-renew and give rise to a much smaller differentiating daughter cell ^67^. We asked whether this could also create a genetic bottleneck that increases variability in mtDNA heteroplasmy levels between daughter cells, similar to the pronounced heteroplasmy differences observed among offspring of the same mother ^24^. A typical example occurs during *Drosophila* neurogenesis in the developing larval brain, where we found NSCs at third instar to be on average ∼12x larger than their differentiating progeny (NSCs:115.9±21.9μm^3^; progeny:9.5±4.2μm^3^; mean±sd) (**Figure 6A**). We conducted mtDNA-smFISH with Dmel and Dyak probe sets on heteroplasmic brains expressing membrane-bound GFP, specifically in NSCs and their direct progeny (*Worniu*-GAL4,UAS-mCD8::GFP) ^68^ and performed GFP co-immunostaining to identify individual NSC lineages. Quantification of mtDNA puncta also showed an average ∼12x reduction in total mtDNA molecules between NSCs and their progeny (NSCs:208.7±68.4; progeny:16.4±7.1; mean±sd) (**Figure 6B**), comparable to the observed cell size reduction. The average mutant heteroplasmy level of NSCs was 66.4±5.7% (n=14 from 3 brains; mean±sd) (**Figure 6C-F**; **Figure 4I**). Interestingly, progeny cells showed significantly greater heteroplasmy variability than NSCs **(Figure 6C,F,G)**, as determined by a likelihood ratio test (LRT) comparing a mixed-effects model assuming different variances for NSCs and progeny to a null model assuming equal variances (LRT=12.446, p=0.0004) (**Figure 6G**). However, using Monte Carlo permutation testing, we found that observed heteroplasmy variance in progeny was not significantly different from that of simulated progeny generated under a random segregation model accounting for mtDNA CN reduction during asymmetric NSC division (p=0.255; **Figure 6H;S9A**). The increased heteroplasmy variance within progeny derived from a single NSC, therefore, is consistent with a model in which the ∼12x reduction in mtDNA CN during asymmetric NSC division acts as a somatic bottleneck, amplifying the effect of random genetic drift on mtDNA segregation.

**Figure 6:**
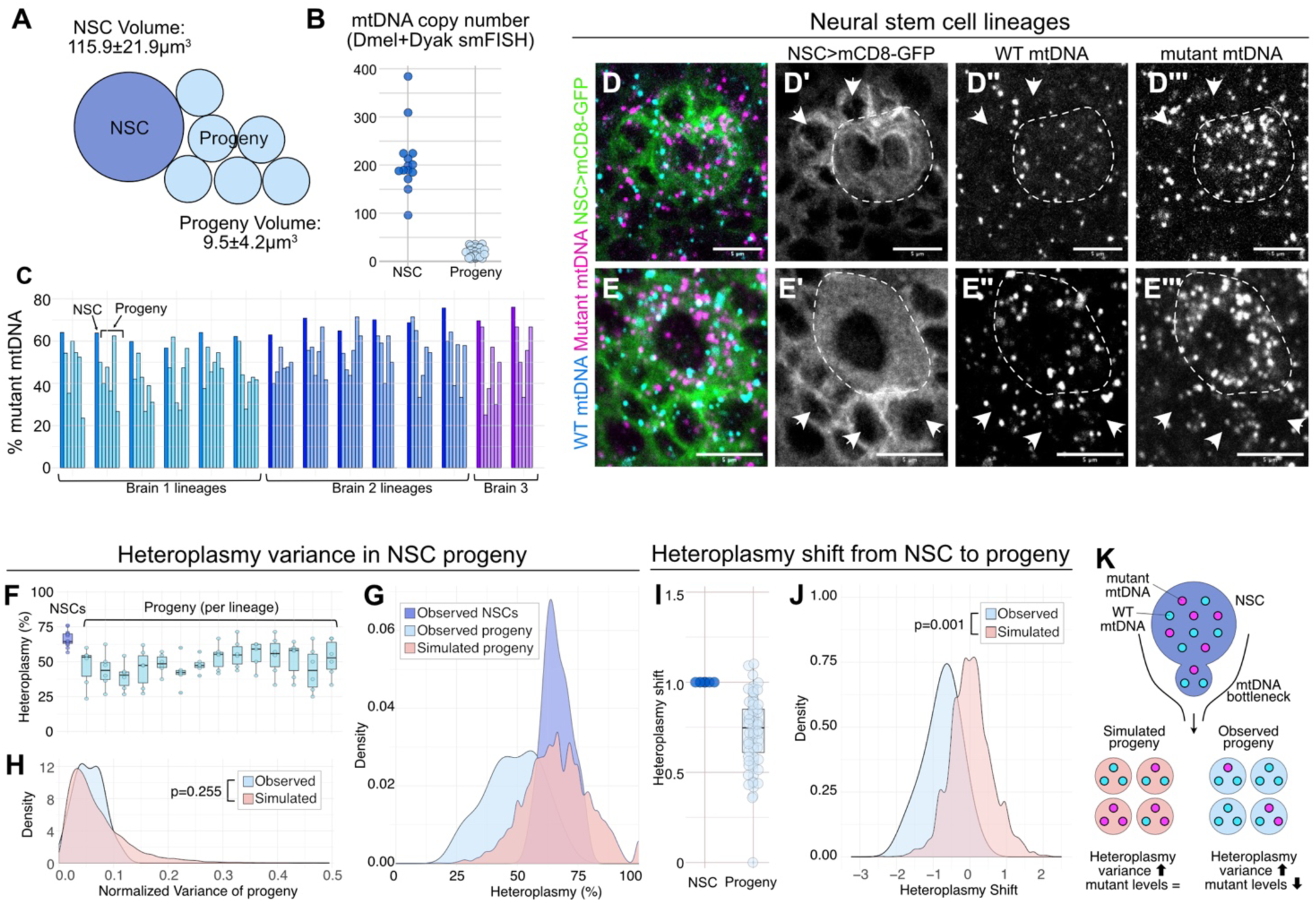
Asymmetric mtDNA segregation through a somatic bottleneck during *Drosophila* neurogenesis. (A) Schematic of *Drosophila* NSC lineage, with NSC volume (dark blue) on average ∼12x larger than their differentiating progeny (light blue). (B) mtDNA CN in NSCs and their progeny using mtDNA-smFISH. (C) Heteroplasmy levels quantified by mtDNA-smFISH in 14 NSC lineages and their youngest-born progeny (identified by Worniu-GAL4 > UAS-mCD8-GFP), across three different brains as indicated. (E,F) Representative images of NSC lineages quantified in (B-D), expressing Worniu-GAL4 > mCD8-GFP (green,D’,E’) and mtDNA-smFISH using probes targeting only *D.yakuba* (WT, cyan,D’’,E’’) or *D. melanogaster* (mutant, magenta, D’’’,E’’’) mtDNA. Outline indicates the NSC and arrows point to smaller progeny. Scale bars are 5μm. (F) Heteroplasmy levels in NSCs pooled across all lineages (dark blue) and their progeny grouped per lineage (light blue). (G,H) Heteroplasmy level distributions (G) and normalized heteroplasmy variance density curves (H) across all lineages for NSCs (dark blue), their progeny cells (light blue) and progeny cells simulated under a random segregation model (red). (I) Heteroplasmy level quantified by mtDNA-smFISH in NSCs and their nearby progeny after normalisation to the heteroplasmy level in each NSC they derived from. (J) Heteroplasmy shift from NSCs to progeny, as observed by mtDNA-smFISH (light blue) or simulated under a random segregation model (red). (K) Model of heteroplasmy segregation through a somatic mtDNA bottleneck during asymmetric NSC division, comparing simulated progeny (red) to observed progeny (light blue) as quantified by mtDNA-smFISH. See also Figure S9.

Interestingly, mutant (Dmel) heteroplasmy levels were almost always lower in the progeny cells than in the NSC they derived from (**Figure 6C-G,I**). A permutation test indeed confirmed that the normalized heteroplasmy shift in observed neurons within each lineage was significantly different from that of simulated neurons (p=0.001; **Figure 6J;S9B**). This indicates that the decrease we observed in the proportion of mutant mtDNA from NSC to progeny, cannot be explained by a random segregation model alone. Asymmetric NSC division and neuronal differentiation must, therefore, involve an asyet-unknown mechanism of purifying selection that favours segregation or selection of functional mtDNA into neuronal progeny (**Figure 6K**).

## Discussion

Heteroplasmy levels for a given mtDNA mutation can show striking cell-to-cell variability, even among the same cell type of a single organ. They also change dynamically over time and thereby impact the pathogenesis of several common and rare inherited neurodegenerative disorders ^69^. Our understanding of the molecular mechanisms that drive or allow cell-type specific accumulation of mtDNA mutations remains limited ^8^, largely due to the inability to efficiently visualize mtDNA sequences in specific cells within their tissue context, and combine this with immunostaining and genetic manipulations. Here, we describe a robust method for visualizing mtDNA heteroplasmy in whole-mount *Drosophila* tissue and in human cells, using sequence-specific smFISH of mtDNA combined with HCR for enzyme-free signal amplification. mtDNA-smFISH enables efficient detection of mtDNA in various *Drosophila* tissues, including the brain, salivary gland, wing discs, muscles, and ovaries. We show that it provides high detection efficacy (>90%) comparing mtDNA-smFISH-based measurements of single-cell mtDNA content with single-cell ddPCR of mtDNA. Specificity was assessed in cell culture and whole-mount tissues by applying mtDNA-smFISH to comparable tissues or cells from other species (human, *D. melanogaster* or *D. yakuba*), and upon expression of a mitochondrial-targeted restriction enzyme to selectively degrade WT mtDNA.

Using this method, we visualized and quantified mtDNA CN and heteroplasmy levels *in vivo* in various developing *Drosophila* tissues, focusing on the female germline and the developing brain, two tissues with key importance for mitochondrial disease inheritance and presentation ^70^. In heteroplasmic *Drosophila* developing brains, we noticed a striking cell-to-cell variability in heteroplasmy levels.

This was evident among individual NSCs in the same brain, in line with single-NSC heteroplasmy measurements by pyrosequencing. However, cell-to-cell variability was significantly higher between postmitotic progeny, even when derived from a single NSC. NSCs undergo asymmetric divisions ^67^ to produce ∼12x smaller progeny, accompanied by a proportionate reduction in mtDNA CN from ∼200 to ∼16 mtDNA molecules per cell. This therefore gives rise to a subcellular somatic genetic bottleneck, whereby only a subset of NSC mtDNA molecules segregates into the differentiating progeny, enhancing genetic drift and inter-neuronal heteroplasmy variability (**Figure 6K**). Statistical modelling of this bottleneck confirms that mtDNA CN reduction is the main factor underlying increased heteroplasmy variance in NSC progeny. Many other instances of asymmetric cell division or cell size reductions occur throughout development and tissue regeneration, for example in hair follicles ^34,71^. This is likely to lead to similar increases in heteroplasmy variance and may underlie the accumulation of clonal mtDNA mutations in some neuron cells, as observed in the brain ^72^.

The rapid segregation of mtDNA heteroplasmy which occurs due to a genetic bottleneck can also expose mutant mtDNA to subtle selection mechanisms ^25,64,73,74^. In *Drosophila*, the developing brain in larvae was previously found to show stronger purifying selection against mutant mtDNA than other tissues like wing disc, gut, or the aging adult brain ^16^. Interestingly, we found mutant mtDNA levels to be consistently lower in progeny than in the NSC they directly derived from. This suggests that neurogenesis is accompanied by an as-yet-unknown mechanism of purifying selection, to ensure that new-born neurons carry more functional mtDNA, limiting the potential deleterious impact of increased heteroplasmy variability caused by a somatic bottleneck. Strikingly, we also observed purifying selection at a well-characterised genetic mtDNA bottleneck during the early stages of *Drosophila* oogenesis. This modulates mother-to-offspring mtDNA inheritance, and has been studied extensively before, mostly through indirect measures like qPCR measurements of mtDNA heteroplasmy levels in offspring ^15,35,63,64,75^. Purifying selection in the female germline is thought to occur through a range of mechanisms, in particular autophagy/mitophagy of mitochondria with mutant mtDNA ^35,63^, and preferential protein translation on the surface of mitochondria with WT mtDNA ^64,75^. Selective mitophagy of mitochondria with damaged mtDNA has also been observed in C. elegans ^76,77^ and human cell culture ^78^. However, within each NSC lineage, almost all progeny contained lower heteroplasmy than the NSC they derived from. Since we only quantified heteroplasmy in the most-recent born progeny, localized close to the parent NSC, it seems likely that, already upon NSC division, mtDNA is subject to asymmetric segregation of WT rather than mutant mtDNA. This is reminiscent of previous observations in mammalian cells ^79,80^ and yeast ^81–83^, that stem cells can selectively segregate old/young mitochondria or WT/mutant mtDNA into daughter cells, and could be driven by the intricate molecular machinery that regulates asymmetric apico-basal apportioning of intrinsic NSC/progeny fate determinants in *Drosophila* neuroblasts ^67,84^. It will be interesting to determine which mechanisms act in NSCs and their progeny, whether these are conserved in other somatic tissues and between species, and whether and how recently-described mtDNA quality control mechanisms like nucleoidophagy ^85–87^ may act specifically on mitochondria containing WT or mutated mtDNA.

Together, the new insights gained from this work contribute to our understanding of mtDNA dynamics, and provide a valuable tool for investigating mitochondrial inheritance and evolution in various contexts, with broader implications for understanding human mitochondrial diseases, neurodegeneration and aging.

## Limitations

Minor cross-reactivity was observed of *D. yakuba*-specific probes with *D. melanogaster* mtDNA, which can be addressed by adjusting thresholds for signal quantification. Another limitation is that mtDNA-smFISH cannot differentiate mitochondrial genomes that only differ by a single-nucleotide polymorphism, as several variants or small indels are needed to confer probe specificity. Precisely how much sequence divergence is required remains to be determined. Nonetheless, when applied in the right heteroplasmic context, and combined with immunostaining and targeted genetic perturbations, mtDNA-smFISH constitutes a powerful novel approach for probing the cellular and molecular mechanisms that shape mtDNA heteroplasmy dynamics during development and aging.

## Acknowledgements

We thank all lab members and F. Karam-Teixeira, T. Samuels and W. Staels for helpful discussions and advice; H. Choi and colleagues from Molecular Instruments for generously sharing reagents and advice; C. Moraes for sharing the H2.1 cybrid cell line; R. Chowdhury, A. Harrison from the MRC MBU Imaging Facility for support with imaging; R. Schulte, G. Grondys-Kotarba from the CIMR Flow Cytometry Facility for assistance with cell sorting; A. Glynos, S. Burr, P. Nash for help with pyrosequencing and ddPCR; L. Bozhilova for advice on statistics and modelling. *Drosophila* stocks obtained from the Bloomington *Drosophila* Stock Center (NIH P40OD018537) were used in this study.

This work was supported by a Wellcome Clinical Research Career Development Fellowship (219615/Z/19/Z), an Evelyn Trust Medical Research Grant (21-25), and a UKRI BBSRC Responsive Mode Research Grant (BB/X00256X/1) to JvdA. PFC and JvdA are supported by a Wellcome Discovery Award (226653/Z/22/Z), a Medical Research Council (MRC) award (MC_PC_21046) to establish a National Mouse Genetics Network Cluster in Mitochondrial Diseases (MitoCluster), and core funding from the MRC to the MRC Mitochondrial Biology Unit (MC_UU_00028/7 and MC_UU_00028/8). HM was funded by a Wellcome Sir Henry Dale Fellowship (202269/A/16/Z) and an ERC Starting Grant (803852). BG acknowledges studentship funding from the Cambridge Trust and the China Scholarship Council.

For the purpose of open access, the authors have applied a Creative Commons Attribution (CC BY) licence to any Author Accepted Manuscript version arising from this submission.

## Author contributions

RC, AMHA and JvdA designed and conducted mtDNA-smFISH experiments. BG and HM provided essential reagents and assisted with cell culture and ddPCR. SP assisted with *Drosophila* genetics and sample preparation. AD performed statistical analysis and modelling. MF assisted with human cybrid cell culture and analysis. HM, PFC and JVDA supervised the project and obtained funding. RC and JVDA wrote the paper, and all authors edited and approved the final manuscript.

## Data availability

Further information and requests for resources and reagents should be directed to and will be fulfilled by the lead contact, Jelle van den Ameele.

## Materials and Methods

### Key resources table

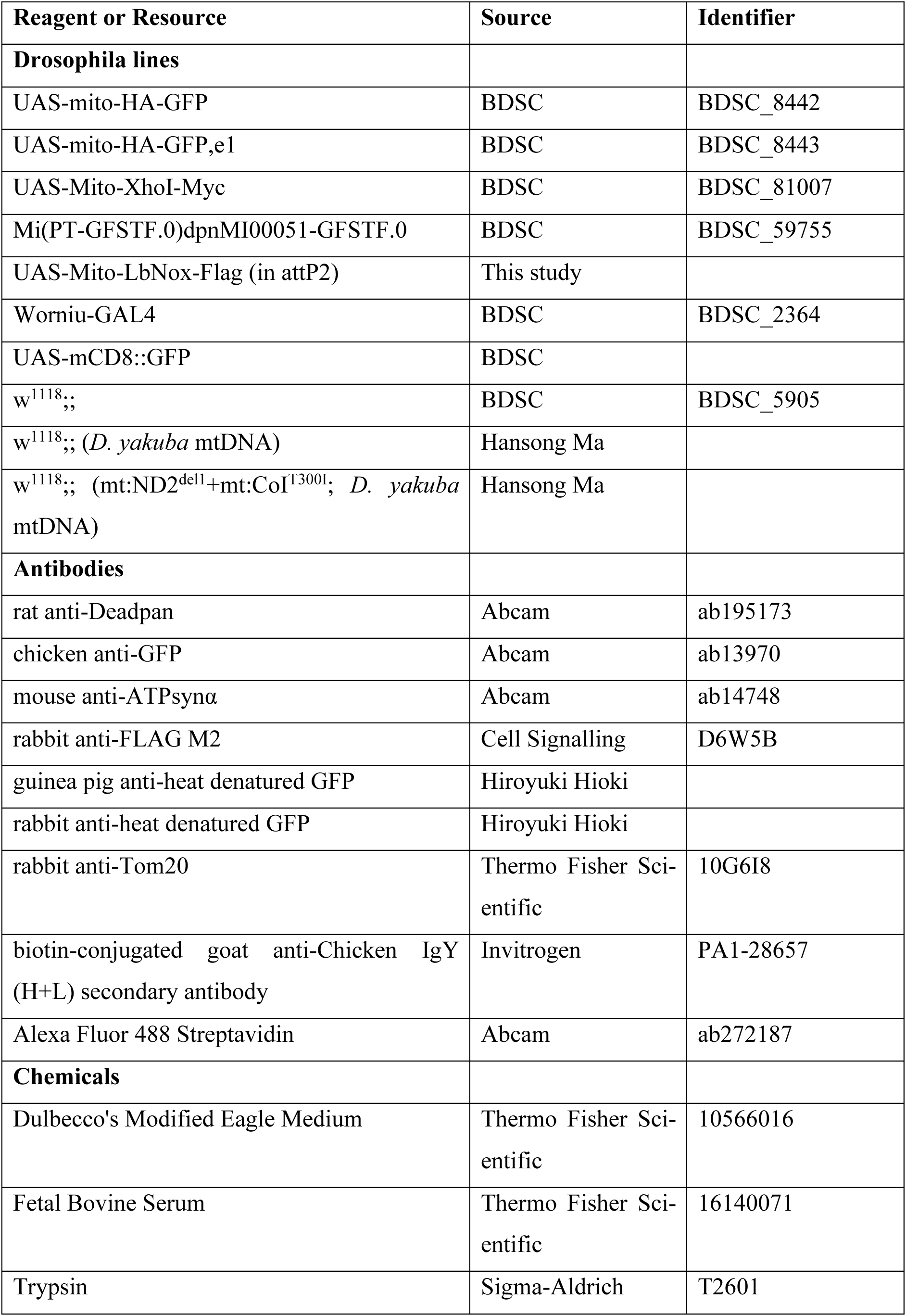
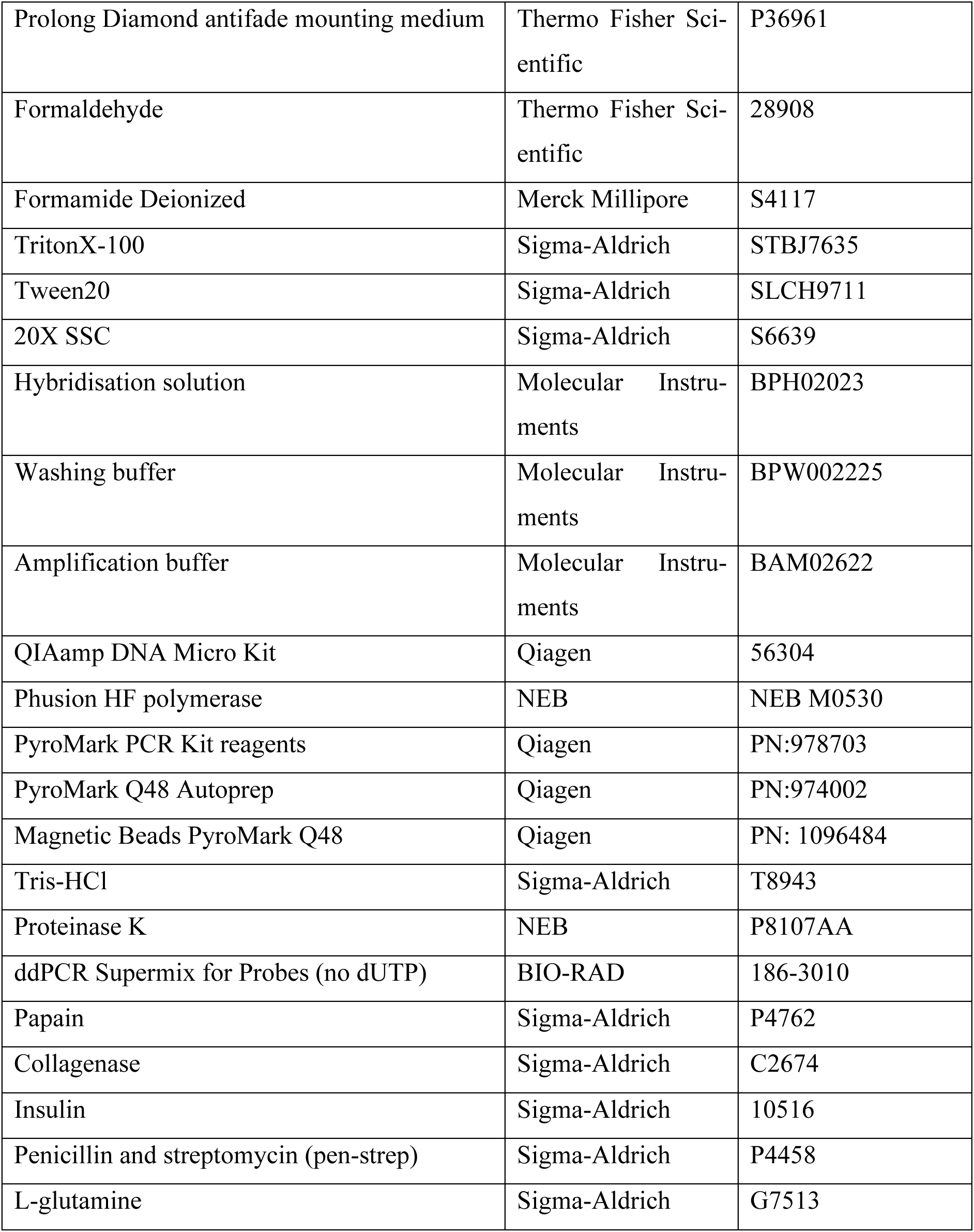

### *Drosophila* strains and husbandry

Homoplasmic Drosophila melanogaster strains were maintained at 25°C, and heteroplasmic strains carrying the temperature sensitive mt.COI^T300I^ mutation at 29°C, unless indicated otherwise. The following stocks were used: UAS-mito-HA-GFP (BDSC_8442); UAS-mito-HA-GFP,e1 (BDSC_8443); UAS-Mito-XhoI-Myc (BDSC_81007); Mi(PT-GFSTF.0)dpnMI00051-GFSTF.0 (Dpn::GFP) (BDSC_59755); UAS-Mito-LbNox-Flag (A. Hynes-Allen and D. Dubal, unpublished). The GAL4-driver used throughout the study was Worniu-GAL4 on II ^53^ (BDSC_2364), either on its own or recombined with UAS-mCD8-GFP on II. Heteroplasmic Drosophila, carrying a combination of wildtype *D. yakuba* mtDNA and mutant *D. melanogaster* mtDNA (mt:ND2^del1^+mt:CoI^T00I^), on a nuclear *D. melanogaster* w^1118^ genomic background, were described previously ^15^. Heteroplasmy was monitored routinely by PCR-restriction digest (see below; **Figure S6**). Apart from the experiments on ovaries, all experiments were done on both male and female larvae.

### Cell culture

*Drosophila melanogaster* S2R+ cells were maintained in Schneider’s *Drosophila* Medium (Thermo Fisher Scientific 21720024) supplemented with 10% fetal bovine serum (FBS) (Thermo Fisher Scientific 16140071), in a humidified incubator at 25°C.

HeLa cells were cultured in Dulbecco’s Modified Eagle Medium (DMEM) (Thermo Fisher Scientific 10566016) with 10% FBS (Thermo Fisher Scientific 16140071) in a closed 10ml dish at 37°C, 4.9% CO2 and 18.0% O2. Medium was refreshed every other day. Upon reaching confluency (every 3-4 days), cells were detached using Trypsin (Sigma-Aldrich T2601) for 3-5 minutes, resuspended in culture medium, spun down at 200xg for 2 minutes at RT, resuspended in 1 ml of fresh medium, and diluted 1:10 in a fresh culture dish.

H2.1 cybrid cells carrying a heteroplasmic mtDNA deletion were obtained from Dr Carlos Moraes ^58^. Cells were cultured in a 5% CO2 and 37°C incubator in DMEM-high glucose-pyruvate (Gibco 41966), supplemented with 10% FBS (Gibco A5256701) and 50 µg/mL uridine (Sigma U3003).

### Antibodies

The following primary antibodies were used: rat anti-Deadpan (1/100, Abcam ab195173); chicken anti-GFP (1/500, Abcam ab13970); mouse anti-ATPsynα (1/500, Abcam ab14748); rabbit anti-FLAG M2 (1/500, Cell Signalling D6W5B); Guinea pig or rabbit anti-heat denatured GFP (1/500, kind gift of Hiroyuki Hioki ^88^); rabbit anti-Tom20 (1/500, Thermo Fisher Scientific 10G6I8). Secondary antibodies for conventional immunostaining were Alexa Fluor-conjugated secondary antibodies raised in goat (1/500, Life Technologies). For biotin-labelled immunostaining, we used biotin-conjugated goat anti-Chicken IgY (H+L) secondary antibody (1/500, Invitrogen PA1-28657) and Alexa Fluor 488 Streptavidin (1/250, Abcam ab272187).

### mtDNA-smFISH probes

mtDNA-smFISH probes for third-generation FISH-HCR ^43,45^ were either ordered from Molecular Instruments, or designed in-house and ordered as desalted unmodified oligonucleotides (Sigma). smFISH performance in tissues was optimal with probes ordered from Molecular Instruments (data not shown). Probes were designed against a target sequence of 52 base pairs, with two halves designed against 25 base pairs on either side, separated by two base pairs. Initiator and spacer sequences for each set of amplifiers were as in ^43,46^. Oligonucleotides were then assembled as follows: (probe half 1) Initiator1 - I1spacer - Reverse complement of 5’side of target sequence; (probe half 2) reverse complement of 3’side of target sequence - I2spacer - Initiator 2. Target sequences, and probe sequences for in-house designed probes are shown in **Table S1**.

### Whole-mount mtDNA-smFISH

Larval tissues were dissected in PBS, fixed in 4% formaldehyde (FA) in PBS with 0.3% TritonX-100 (Sigma-Aldrich STBJ7635) (0.3%PBSTx) for 30 minutes at RT, and washed for 3 × 5 minutes in 0.1% PBSTx at RT. Fixed tissue underwent dehydration through consecutive washes in cold PBS with 0.1% Tween20 (Sigma-Aldrich SLCH9711) (PBSTw) with 25%, 50%, 75%, and 100% methanol for 10 minutes each at 4°C. Tissue can be stored at −20°C at this stage, until *in situ* hybridisation (ISH). Prior to ISH, tissues were rehydrated through consecutive washes in cold PBSTw with 100%, 75%, 50% and 25% methanol for 10 minutes each, and a final wash in PBSTw. Denaturation solution was 70% Formamide (Merck Millipore S4117), 2x SSC (Sigma-Aldrich S6639) in H_2_O. Tissues were incubated in 600μl denaturation solution at 90°C in a shaking heating block at 300rpm for 10 minutes. Probes were stored at 1 μM stock solution (i.e. 1 μM for each probe half) at −20°C, defrosted prior to ISH, and 0.8μl was added to 200μl hybridisation solution (Molecular Instruments BPH02023; stored at - 20 °C), for a final hybridisation concentration of 4nM for each probe-half. Final total hybridisation concentration therefore depends on the number of probes in each probe set. For smaller probe sets (i.e. Dmel and Dyak specific probe sets targeting 7 and 5 sequences respectively), concentration was increased to up to 40nM, to improve signal intensity. Probes were heated in hybridisation solution to 95°C for 2 minutes. Denaturation solution was removed from the tissues and replaced by the heated hybridisation solution with probes. Heating block temperature was adjusted to 37°C, to rapidly but gradually cool the tissues with probes, and then incubated overnight at 37°C in a shaking heat block. In the meantime, washing buffer (Molecular Instruments BPW002225; stored at −20°C) was placed in a 37°C incubator to pre-warm. The following day (day 2), tissues were washed 4 x 15 minutes with warmed probe wash buffer at 37°C (for the first wash, wash buffer was added to the sample directly to make tissues sink to the bottom of the vial), followed by a 5 minutes wash in 50% probe wash buffer and 50% 5xSSC with 0.1% Tween20 (SSCT) at RT and two final 5 minutes washes with SSCT at RT. HCR amplifiers were defrosted in the dark, and annealed by placing 2μL (of 3μM stock for 6pmol final concentration per sample) of each hairpin in separate tubes, heated to 95°C for 1.5 minutes and cooled at RT in the dark. Next, all HCR hairpins were combined and added to amplification buffer (Molecular Instruments BAM02622; stored at 4 °C; 100µL per sample), if necessary, with nuclear counterstain DAPI (4’,6-diamidino-2-phenylindole) at 1:500, added to the tissue sample and incubated in the dark overnight at RT. The final day (day 3) consists of washes with SSCT of 2 x 5 minutes, 2 x 30 minutes, and 1 x 5 minutes, followed by 3 x 10 minutes washes with 0.1%PBSTx and 2 x 10 minutes with PBS. Tissues were mounted in ProLong Diamond antifade mounting medium (Thermo Fisher Scientific P36961). Slides were stored at 4℃ and imaged after 24 hours.

### mtDNA-smFISH on cultured cells

HeLa or S2R+ cells were cultured in IBIDI 12-Well Chambers (Thistle Scientific IB-81201) in 250µL of medium, the maximum capacity for the wells. Cells were fixed using 4% formaldehyde (FA) in PBS containing 0.1% PBSTx, washed in PBSTx, and subjected to dehydration and rehydration steps as per the smFISH protocol. For heating during the pre-hybridisation step, the IBIDI plate was placed on a flat surface of an inverted heating block underneath a polystyrene cover. For probe hybridisation, the IBIDI wells were placed in a wet chamber to prevent evaporation. This chamber was created by placing pre-warmed, wet tissue paper at the bottom of an empty pipette tips box, with the IBIDI wells on top. The box was maintained at 37°C in an incubator. Subsequent wash steps, and antibody staining were essentially as for the whole-mount smFISH-HCR protocol, as described above.

### Incorporating immunostaining with mtDNA-smFISH

For conventional antibody staining of epitopes not affected by heat denaturation (e.g. anti-Flag immunostaining), or with antibodies against heat denatured GFP ^88^, immunostaining was conducted after the smFISH protocol. In the final washes, 5xSSCT was replaced by 0.3%PBSTx through serial washing with 75%, 50% and 25% 5xSSCT in PBST for 5 minutes each, and 3 x 5 minutes in PBST. For immunostaining, brains were incubated with primary antibodies in 0.1%PBSTx overnight at 4°C, washed with 0.1%PBSTx, incubated with secondary antibodies in PBST overnight at 4°C and washed with 0.1%PBSTx. Brains were mounted in Prolong Diamond antifade mounting medium (Thermo Fisher Scientific P36961).

For biotinylated antibody staining, primary and secondary biotinylated antibody staining followed by a second fixation step were conducted before the smFISH protocol. After the rehydration step on day 1, tissues were incubated with anti-GFP chicken primary antibody (Abcam ab13970; 1:500) in 0.1% PBSTx overnight at 4°C. The following day (day 2), tissues were washed 3 x 10 minutes with 0.1% PBSTx, and incubated overnight with goat biotinylated anti-chicken IgY (H+L) secondary antibody (Invitrogen, PA1-28657; 1:500). The next day (day 3), tissues were washed 2 x 5 minutes with 0.1% PBSTx, followed by a second fixation in 4% FA in 0.1% PBSTx for 20 minutes at RT, and washed 2 x 5 minutes in PBSTw, to next continue with the mtDNA-smFISH denaturation steps and protocol. In the final washes of the smFISH protocol, the tissues were processed as described above for conventional immunostaining, but after the final wash in 0.1%PBSTx, incubated overnight with streptavidin Alexa Fluor 488 conjugate at 1:100 (Jackson Immunoresearch 016-540-084). The final day, 3 x 10 minutes washes were done with 0.1%PBSTx, and 2 x 10 in PBS prior to mounting in Prolong Diamond antifade mounting medium (Thermo Fisher Scientific P36961).

### *Drosophila* larval brain dissociation and fluorescence-activated cell sorting

Dissociation of *Drosophila* larval brain tissue was done essentially as described previously ^89^. About 50 L2/L3 larval brains were dissected in ice-cold PBS and transferred to an Eppendorf low binding microcentrifuge tube with 100µl of ice-cold Rinaldini’s solution (800mg NaCl, 20mg KCl, 5mg NaH_2_PO_4_, 100mg NaHCO_3_, 100mg glucose in 100ml distilled H_2_O; filtered through Millipore Steriflip (pluriSelect); 10x can be stored at 4°C). Brains were centrifuged at 300g for 3 minutes at 4°C and the supernatant replaced with dissociation solution (445μl Schneider’s medium, 50 papain (Sigma-Aldrich P4762) and 5μl collagenase (1mg/ml, Sigma-Aldrich C2674) (Schneider’s culture medium consists of 2.5ml fetal bovine serum (Thermo Fisher Scientific 16140071), 50μl insulin (Sigma-Aldrich 10516), 500μl pen-strep (Sigma-Aldrich P4458), 2.5ml L-glutamine (Sigma-Aldrich G7513) in 18.925ml Schneider’s medium (Thermo Fisher Scientific 21720024) and filtered through Millipore Steriflip (pluriSelect). Tissues were incubated at 30°C in a shaking heat block for one hour, with gentle pipetting every 15 minutes. Next, dissociated cells were centrifuged at 300xg for 3 minutes at 4°C, washed with 1ml ice-cold Rinaldini’s solution and centrifuged at 300xg for 3 minutes at 4°C. Subsequently, 200µl Schneider’s culture medium was added, cells resuspended by 10-20 times pipetting with a 200µl pipette tip, centrifuged again at 300xg for 3 minutes at 4°C and finally resuspended in 400μl PBS with 0.04%BSA. The cell suspension was filtered through a 10µm pluriStrainer (Cambridge Bioscience 43-50010-03), after wetting the filter with 200μl PBS with 0.04%BSA. Fluorescence-activated cell sorting (FACS) was carried out on a Melody^Tm^ Cell sorter (BD Biosciences), and data analysis was conducted on FlowJo software (BD Biosciences). The cell suspension after tissue dissociation was transferred to a 5ml 12×75mm style (Falcon) polystyrene round bottom tube with a cell strainer cap, and kept on ice before FACS. FACS settings were optimised to keep living and non-permeabilised cells (DAPI-negative and DRAGQ5) (**Figure S3**) and single or specific numbers of Dpn::GFP-positive cells were collected in a 96-well PCR plate. Plates were immediately stored at −80°C for future downstream processing.

### Droplet Digital PCR (ddPCR)

ddPCR was conducted as described previously ^90^. For mtDNA CN measurements with ddPCR on sorted NSCs, cells were processed in a 96-well plate, either as single cells, or as sets of 10, 20, 50 or 100 cells. Cells were lysed in 2.5µL lysis buffer consisting of 50mM Tris-HCl (Sigma-Aldrich

Trizma T8943) at pH 8.3, TWEEN-20 (Sigma-Aldrich SLCH9711), and 200µg/mL Proteinase K (New England Biolabs P8107AA). Plates were centrifuged at 1,000 x g for one minute at 4°C, incubated at 37°C for 30 minutes in a thermocycler with the lid heated at 105°C, and Proteinase K was next heat inactivated for 15 minutes at 80°C. 7.5µL of nuclease-free water was added to each sample to a final volume of 10µL, and plates were centrifuged at 1,000 x g for one minute at 4°C. The cell lysate was stored at −20°C before proceeding with ddPCR if needed. ddPCR Master mix for 1 reaction consists of 12.5µL 2x ddPCR Supermix for probes (BIO-RAD 186-3010), 1.25µL of 20x VIC, 1.25µL of 20x FAM probes and 7µL of sterile water, to a final volume of 22µL, and vortexed after preparation. In this protocol, VIC targets both Dmel and Dyak mtDNA, while FAM targets only Dmel mtDNA (**Figure S4A**). PCR primer sequences were (all 5’-3’): Dyak+Dmel PCR Forward, AACAGACTTAAAATTTGAACGGCTACAC; Dyak+Dmel PCR Reverse, GGAGAGTTCATA-TCGATAAAAAAGATTGCG; Dmel PCR Forward, CTTCCAGTACTAGCAGGAGCTATT; Dmel PCR Reverse, CCTCCTCCCGCTGGGT. Probe sequences were (all 5’-3’): D. yakuba and D. melanogaster mtDNA (VIC), TATATCTTAATCCAACATCGAG; D. melanogaster mtDNA (FAM), ATCGAAATTTAAATACATCATTTTTT. 22µL master mix was added to each well of a new 96-well plate, and 3µL of DNA from the lysed cells, or nuclease-free water to the non-template control wells to bring total volume to 25µL. Plates were sealed, centrifuged at 1,000 x g for 1 minute at 4°C and transferred to the droplet generator. Once droplets were formed, PCR was performed for 40 cycles with denaturation at 94°C for 30 seconds and annealing and extension at 58°C for 1 minute. Finally, the 96-well plate is placed in the droplet reader for analysis. mtDNA CN was calculated as follows: mtDNA target concentration/µL x 25 x 1/fraction of total lysate input. FAM probe was specific for Dmel mtDNA (**Figure S4B**), and mtDNA CN calculations of Dmel mtDNA across a range of mtDNA concentrations were similar for FAM and VIC probes (**Figure S4C,D**). mtDNA CN scaled according to cell numbers (**Figure S4C-H**), further validating the assay and allowing to plot all percell values from these assays together. The assay variability was higher among single cells than among sets containing a group of cells (10, 20, 50 or 100), possibly due to cell doublets caused by inefficient FACS sorting (**Figure 2D;S4F,H**).

### PCR restriction digest to determine heteroplasmy

We developed a straightforward genotyping approach, to monitor Dmel and Dyak heteroplasmy levels at low cost, taking advantage of an NdeI restriction site in the Dyak mtDNA (**Figure S8A**), with NdeI retaining 100% in most PCR buffers. Genomic DNA was extracted from whole flies or dissected with QIAamp DNA Micro Kit (QIAGEN 56304). PCR was done with Phusion HF polymerase (New England Biolabs, ID: NEB M0530), for 30 cycles at melting temperature of 45°C, using the following primers common to both Dmel and Dyak mtDNA: Forward 5’-TTAGACCAAATTTATTGGGAGACCC-3’ and Reverse 5’-GGAATTCCTCAACCTTTTT-GTGATGC-3’, resulting in 1283bp Dmel and 1288bp Dyak fragments. NdeI (NEB R3131) digests the Dyak but not Dmel mtDNA, resulting in 2 smaller fragments (425 and 859bp) (**Figure S8B**). Heteroplasmy levels were inferred by measuring intensity of the smaller Dyak bands vs the larger Dmel band (**Figure S8B**), either after imaging of the agarose gel, or on a Tapestation (Agilent).

### Bulk and single-cell pyrosequencing of Drosophila mtDNA variants

In order to accurately measure heteroplasmy levels with high sensitivity in single cells, we developed a pyrosequencing approach ^91,92^ to distinguish Dyak from Dmel mtDNA (**Figure S8C-F**). Pyrosequencing was done using the QIAGEN protocol, with PyroMark PCR Kit reagents (QIAGEN PN:978703), and primer design according to the PyroMark Assay Design SW 2.0 quick start guide. We determined optimal annealing temperatures for several primer pairs to allow efficient PCR amplification of the mtDNA target region. Validation was conducted through pyrosequencing of homoplasmic Dyak or Dmel animals on a standard curve of mixed PCR products (**Figure S8E,F**) and by comparing two different sets of primers (ID8 and ID9), together confirming the accuracy of heteroplasmy detection. Pyrosequencing primers (all 5’-3’) were Forward-8 TTTAATATTTGGTCCTTTCGTACT and Reverse-8 AGCCAGGTTGGTTTCTATCTTTAA, with sequencing primer GGTTTCTATCTTTAAAAAAT; Forward-9 GCTACCTTTGCACAGTCAAAATAC and Reverse-9 GTTTAAATAAAGAATTCGGCAAAA, with sequencing primer TTTGTTAAACAGGCGA. Pyrosequencing of heteroplasmic animals yielded reproducible results (**Figure S8E,F**), with values in line with our first PCR- and digestion-based method (data not shown), and with an independent, previously validated qPCR-based method ^61^ (**Figure S8E,F**).

Single-cell pyrosequencing was conducted on sorted NSCs (**Figure S3**). Before processing, the 96-well plate was spun down at maximum speed (2204xg) at 4℃ for 20 minutes, and samples were processed in a dedicated PCR hood with UV treated equipment. PyroMark PCR Master mix was pipetted straight onto the single-cell 96-well PCR plate, spun down for 1 minute at 300xg and PCR was for 45 cycles with melting temperature of 62°C. 2μl of the PCR was verified on a 0.9% agarose gel, and PyroMark Q48 Autoprep (Qiagen) was used to conduct pyrosequencing of 10μl sample in a 48-well disc pyrosequencing plate with 3μl of Magnetic Beads PyroMark Q48 (PN:1096484).

### Confocal and Airyscan laser scanning microscopy, image analysis

Fluorescent images were acquired using a Zeiss LSM880 confocal microscope, equipped with Fast AiryScan, with a 63x oil immersion objective, as 12- or 16-bit images. All images are single sections, unless indicated otherwise. For quantifications, slice thickness was ∼0.8μm, with a 1/3 step size, ensuring complete NSCs or germaria were imaged and reconstructed. Images were processed for brightness and contrast using ImageJ. For automated quantification of heteroplasmy, stacks were cropped to individual NSCs, and MorphoLibJ ^93^ was used for segmentation of the Worniu-GAL4,mCD8::GFP IHC signal, to create a mask for downstream quantification with FishQuant ^94^. Segmentation of germaria was performed manually in the X-Y axis, maintaining the entire imaging Z-stack. mtDNA-smFISH puncta were quantified using FishQuant ^94^, using either the germaria or the NSC mask as input. Single NSCs were studied to generate a detection setting, which was then applied as batch mode for other replicates. The filters were modified manually to target the mtDNA spots for each experiment. When NSCs were difficult to segment, we instead used the CellCounter plugin in FiJi ^95^ to count mtDNA puncta manually. Occasionally, gaussian blur was applied prior to quantification. For signal and background quantification, similar areas of the larval ventral nerve cord were selected in FiJi, and the raw data saved as .tiff files, for export to Matlab (The MathWorks Inc.), where the number of pixels at a given intensity value in 256 bins was quantified. Graphs are shown at a log scale to enhance the differences in the lower values. Graphs were made in R, and figures were compiled in Affinity Designer.

### Statistical analysis and modelling

For each experiment, the number of brains analysed ranged from 1-10. Each experiment where statistical analysis was performed included at least two biological replicates. Prism (GraphPad Software) and Excel (Microsoft) were used to perform statistical tests. Shapiro-Wilk test was used to verify normality, and F-test to compare variance. Wilcoxon Signal Rank test or paired t-tests were performed to determine significance. For the mixed-effects modelling analysis, cell type was specified as a fixed effect, and lineage was included as a random effect to account for baseline heteroplasmy differences between lineages using the ‘lm’ function from the R package ‘nlm’. A likelihood ratio test comparing a model with varying variances for NSCs and progeny to a null model assuming equal variances was performed to assess the significance of variance differences between NSCs and progeny. To test whether the observed heteroplasmy variance in the progeny cells could arise from a purely random segregation process during budding, we simulated heteroplasmy levels in progeny based on the total mtDNA CN and heteroplasmy levels of their parent NSC. Specifically, the mtDNA molecules in each progeny cell were sampled from their respective parent NSC using hypergeometric sampling, mimicking the stochastic nature of mtDNA segregation. We calculated normalized heteroplasmy variance for a cell-population with heteroplasmies *h* = (*h*_1_, *h*_2_,…, *h_n_*) as: *V*′(*h*) = *var*(*h*) / *h̄* (1 − *h̄*) as in^4^. This allows calculation of the sample variance independent of the mean heteroplasmy *h̄*. A total of 1000 simulations per progeny were performed providing a null distribution for normalized heteroplasmy variation under a random segregation model, against which the observed normalized heteroplasmy variation was compared using Monte Carlo permutation tests to assess significance at a significance level of α=0.05. To test whether the observed heteroplasmy shifts between NSCs and their progeny could result from random segregation, we calculated the normalized heteroplasmy shift (Δ′*h*) between the NSC and their progeny as in ^96^:

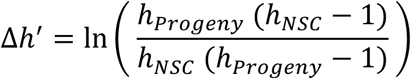

We then compared the observed normalized heteroplasmy shift to a null distribution generated by Monte Carlo permutation testing (1000 permutations), using a significance level of α=0.05.

## Supplementary figure legends

**Supplementary Figure 1:**
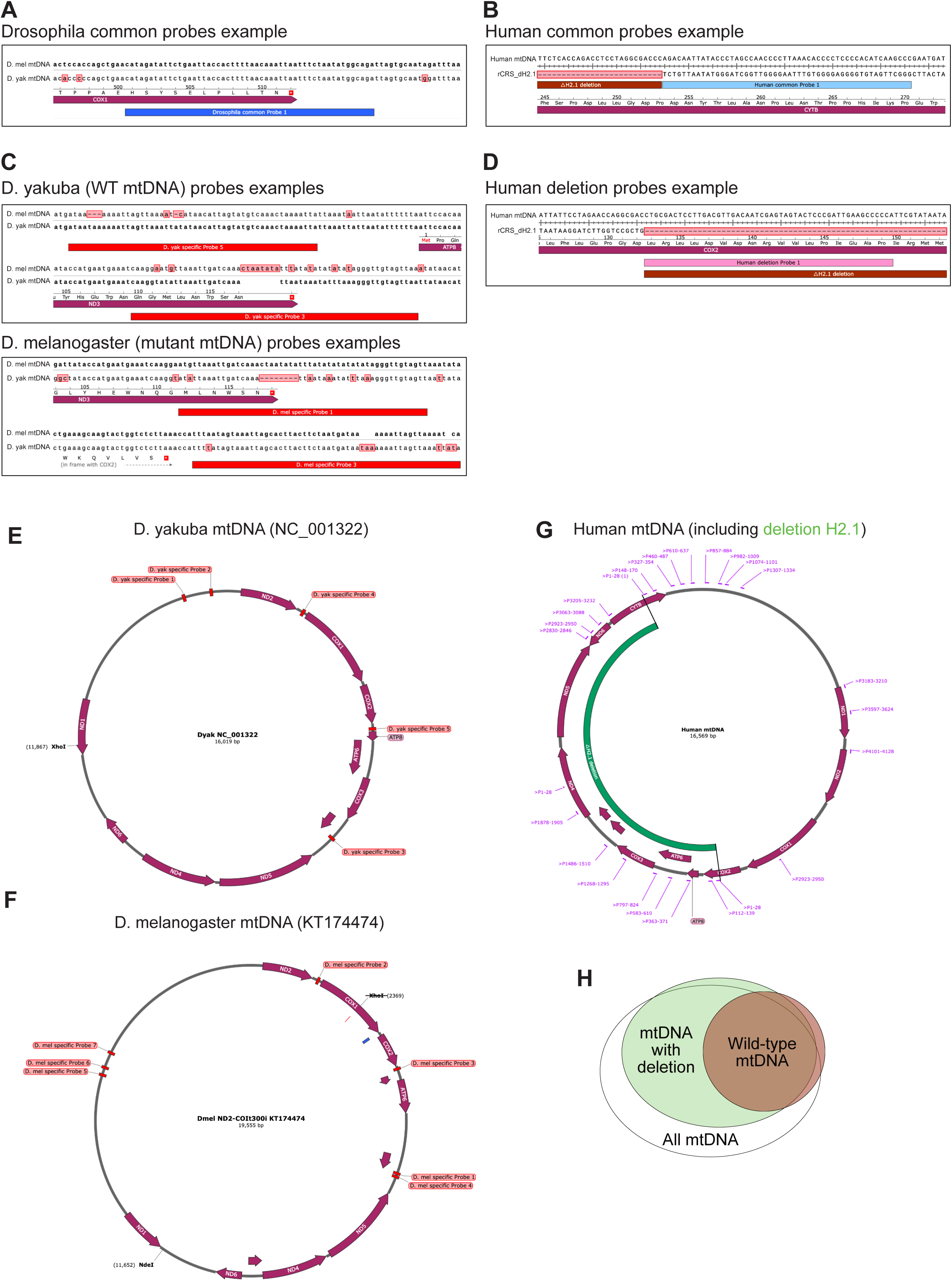
Probe Design for mtDNA-smFISH of *Drosophila* and Human mtDNA. (A-D) Representative examples of target sequences designed for *Drosophila* mtDNA (A,B) common to both *D. melanogaster* and *D. yakuba* (A), specifically for *D. yakuba* (WT, B) or *D. melanogaster* (mutant, B) mtDNA, or for human mtDNA (B,D) common to WT and deleted mtDNA (B) or only recognising the WT mtDNA (D). (E-G) Schematic of *D. yakuba* (E), and *D. melanogaster* (F) mtDNA and human mtDNA (G) with protein-coding genes in purple and probe target sequences in bright red or pink. Human deleted sequences is indicated in green (G). (H) Schematic of how mtDNA-smFISH can distinguish between WT or deleted mtDNA, with deleted molecules only targeted by common mtDNA probes (red) and WT molecules also targeted by deletion-specific probes (green).

**Supplementary Figure 2:**
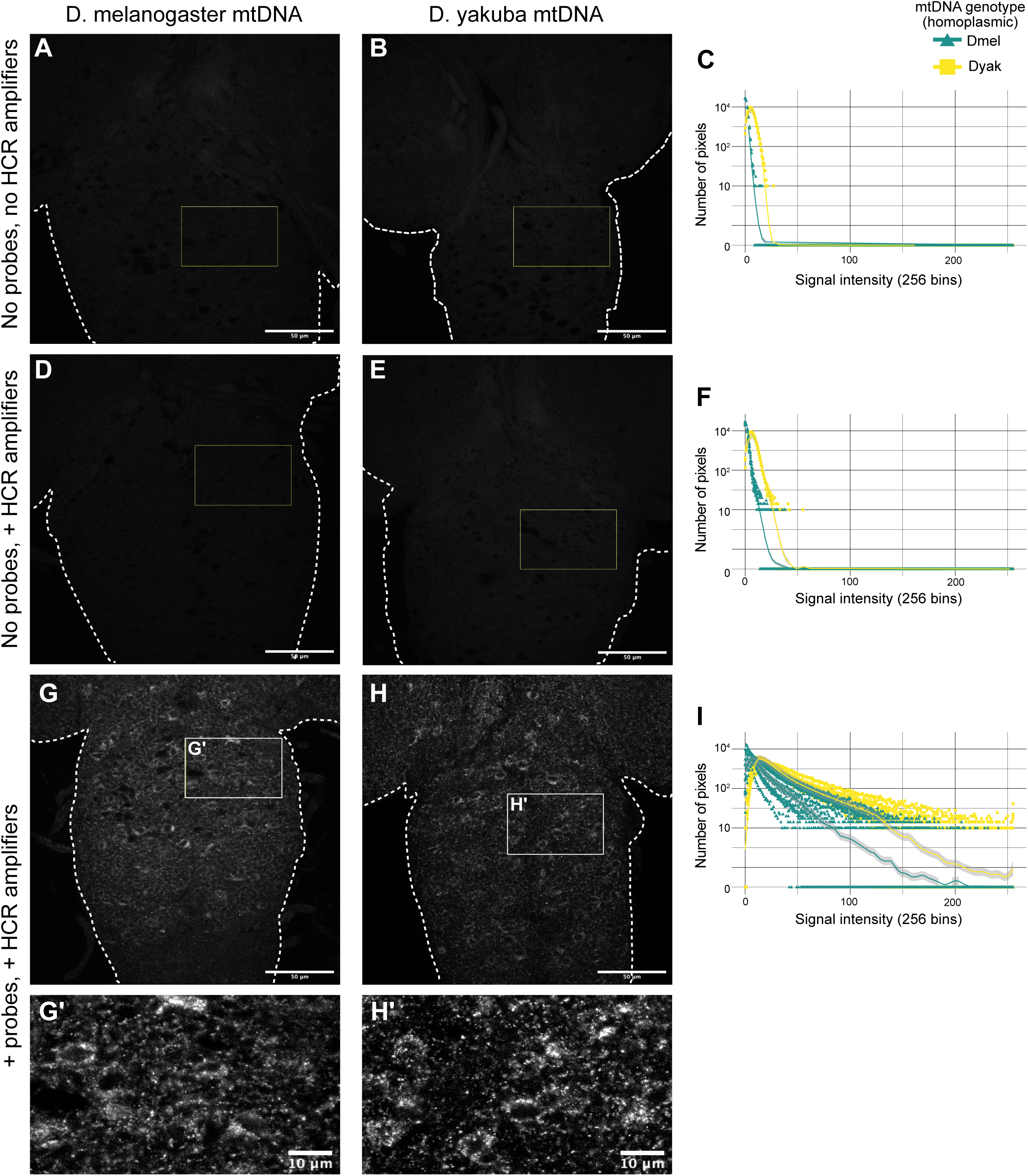
mtDNA-smFISH with probes common to *D. melanogaster* and *D. yakuba* mtDNA. (A-I) mtDNA-smFISH images (A-H) and signal quantification (C,F,I) with probes common to both *D. melanogaster* and *D. yakuba* mtDNA in brains of larvae homoplasmic for *D. melanogaster* (A,D,G) or *D. yakuba* (B,E,H) mtDNA, either without probes or amplifiers (A,B,C), without probes (D,E,F), or with both probes and amplifiers added (G,H,I). (G’,H’) are high magnifications of areas boxed in G,H. Scale bars are 50μm (A,B,D,E,G,H) or 10μm (G’,H’).

**Supplementary Figure 3:**
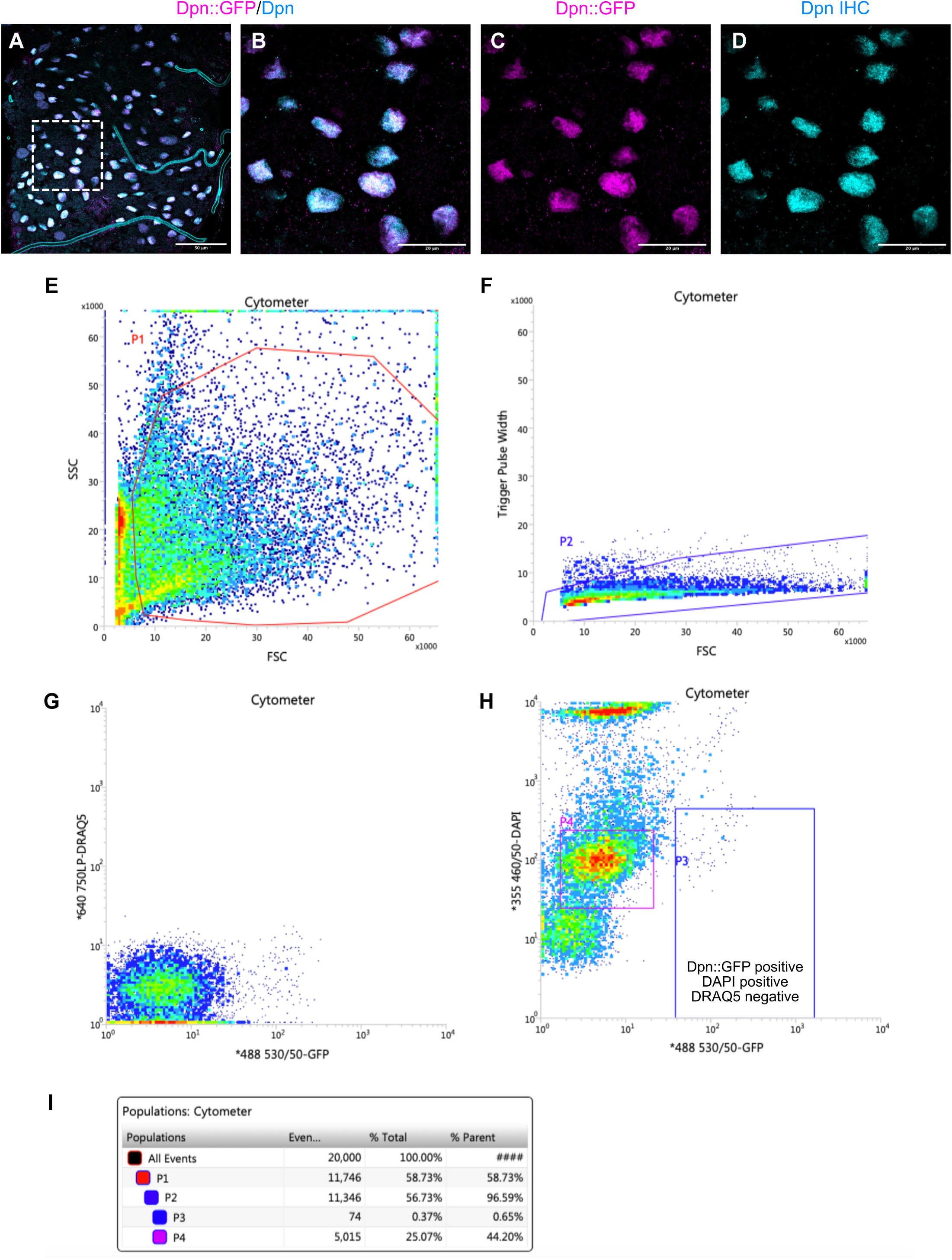
FACS-Based Isolation of *Drosophila* NSCs. (A-D) Immunostaining of Dpn (cyan) and GFP (magenta) of third instar larval brain from transgenic *Drosophila* with GFP fused to the NSC-marker Dpn (Dpn::GFP). (E-I) Representative FACS plots of dissociated third instar larval brains from Dpn::GFP *Drosophila*, with Draq5 and DAPI used to assess viability (G,H), resulting in ∼0.37% of all cells retained as NSCs (I).

**Supplementary Figure 4:**
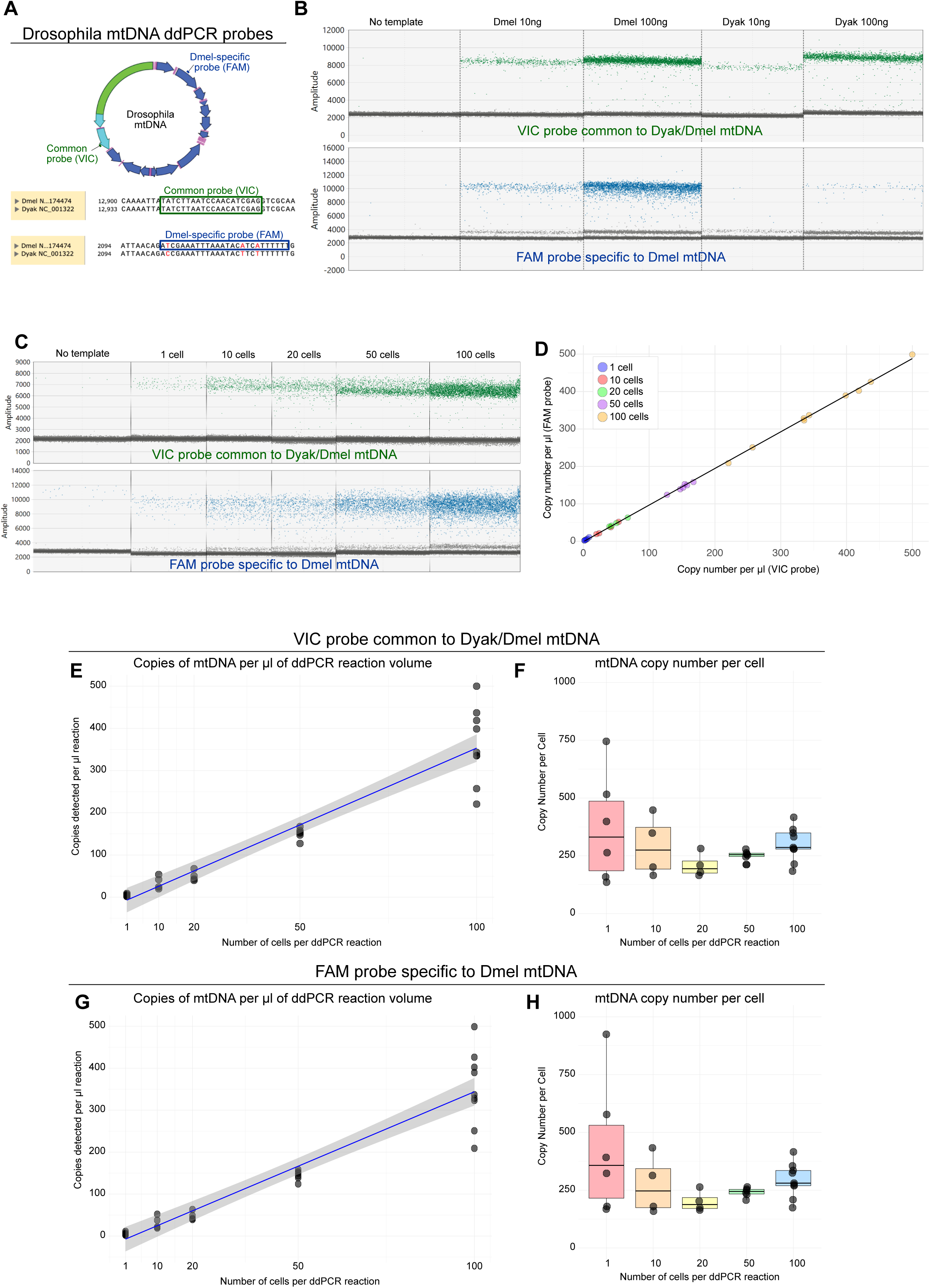
Droplet digital PCR (ddPCR) for *Drosophila* mtDNA. (A) Schematic representation of VIC and FAM probes for *Drosophila* mtDNA. VIC probe was designed to target both *D. melanogaster* and *D. yakuba* mtDNA. FAM probe was specific to *D. melanogaster* mtDNA. (B,C) ddPCR using VIC (green) and FAM (blue) probes using 0, 10 or 100ng of genomic DNA from homoplasmic *D. melanogaster* or *D. yakuba* (B), or on 1, 10, 20, 50 or 100 sorted NSCs (Dpn::GFP) from homoplasmic *D. melanogaster* (C). (D) Correlation between mtDNA CN estimates of VIC and FAM probes on sorted NSCs from homoplasmic *D. melanogaster*. (E-H) mtDNA CN measured using VIC probe (E,F) or FAM probe (G,H) on 1, 10, 20, 50 or 100 sorted NSCs from homoplasmic *D*. *melanogaster*. mtDNA CN increases linearly with increasing cell number (E,G), but larger variability is observed in single-cell samples (F,H).

**Supplementary Figure 5:**
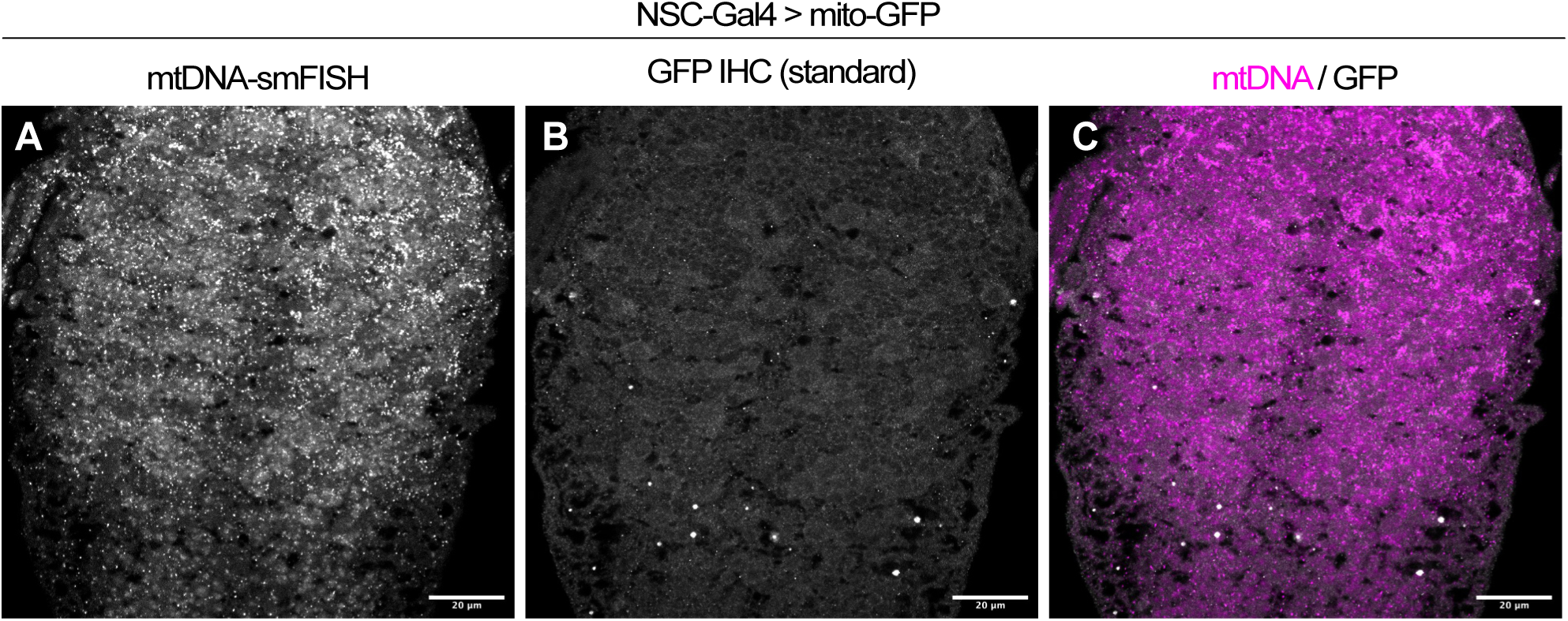
Denaturation prevents conventional GFP immunostaining after mtDNA-smFISH. mtDNA-smFISH (A, magenta) and conventional immunostaining for GFP (Worniu-GAL4 > UAS-mito-GFP) (B, gray) does not effectively stain GFP due to the high denaturation temperature required for mtDNA-smFISH. Scale bars are 50μm.

**Supplementary figure 6:**
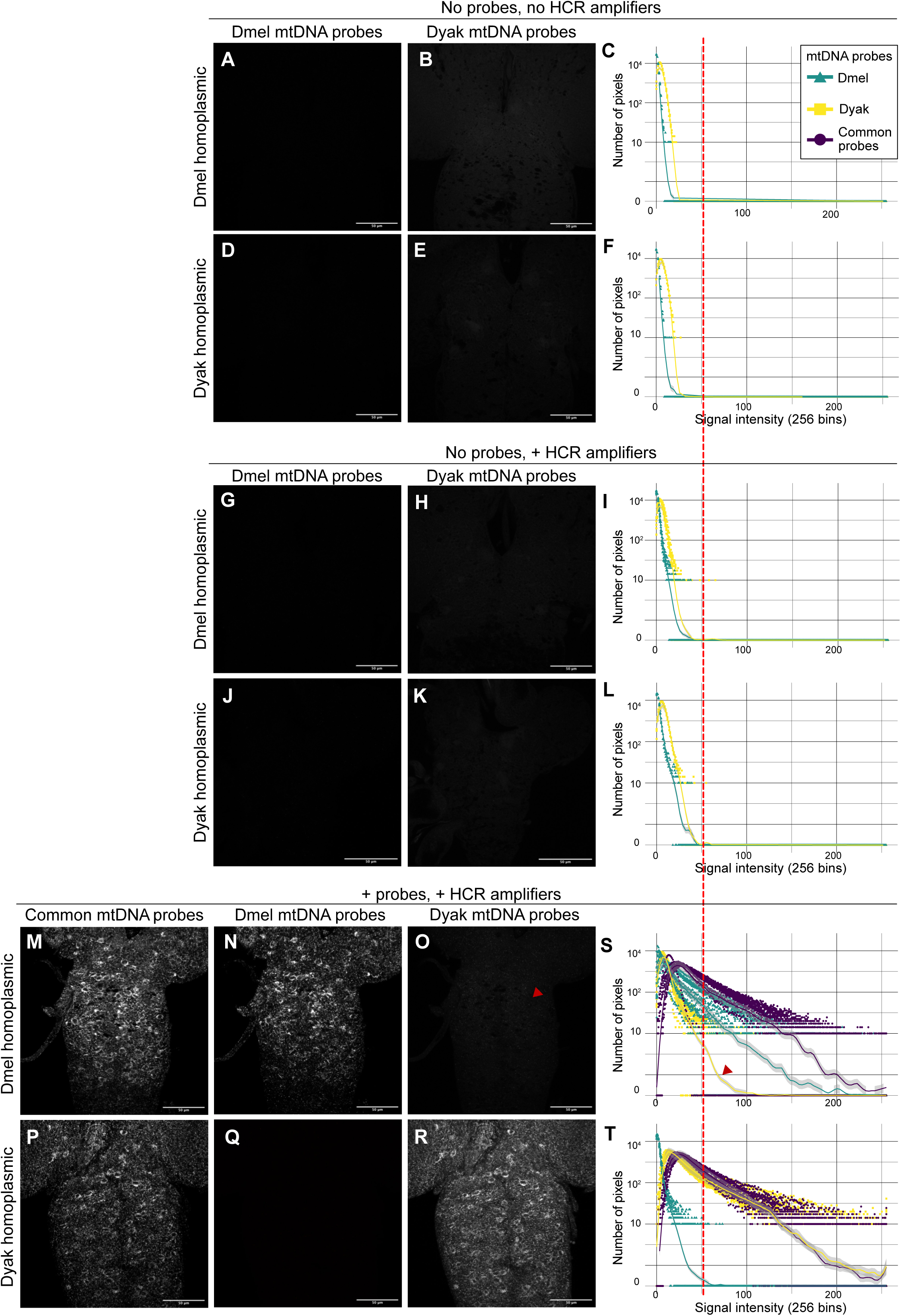
Specificity of *D. melanogaster* or *D. yakuba* mtDNA-smFISH. mtDNA-smFISH images (A-E,G-K,M-R) and signal quantification (C,F,I,L,S,T; mean ± s.e.m.) with probes specific to either *D. melanogaster* (mutant, A,D,G,J,N,Q, green graphs), *D. yakuba* (B,E,H,K,O,R, yellow graphs) or common to both (M,P, purple graphs) mtDNA in brains of *Drosophila* larvae homoplasmic for *D. melanogaster* (A-C,G-I,M-O) or *D. yakuba* (D-F,J-L,P-R) mtDNA, either without probes or amplifiers (A-F), without probes (G-L), or with both probes and amplifiers added (M-R). Scale bars are 50μm. Red arrowheads indicate a low-level non-specific background from the *D.yakuba* probes in *D.melanogaster* homoplasmic (mutant) background. The red line in the graphs separates true signal from background, with a threshold set at 55 bins.

**Supplementary Figure 7:**
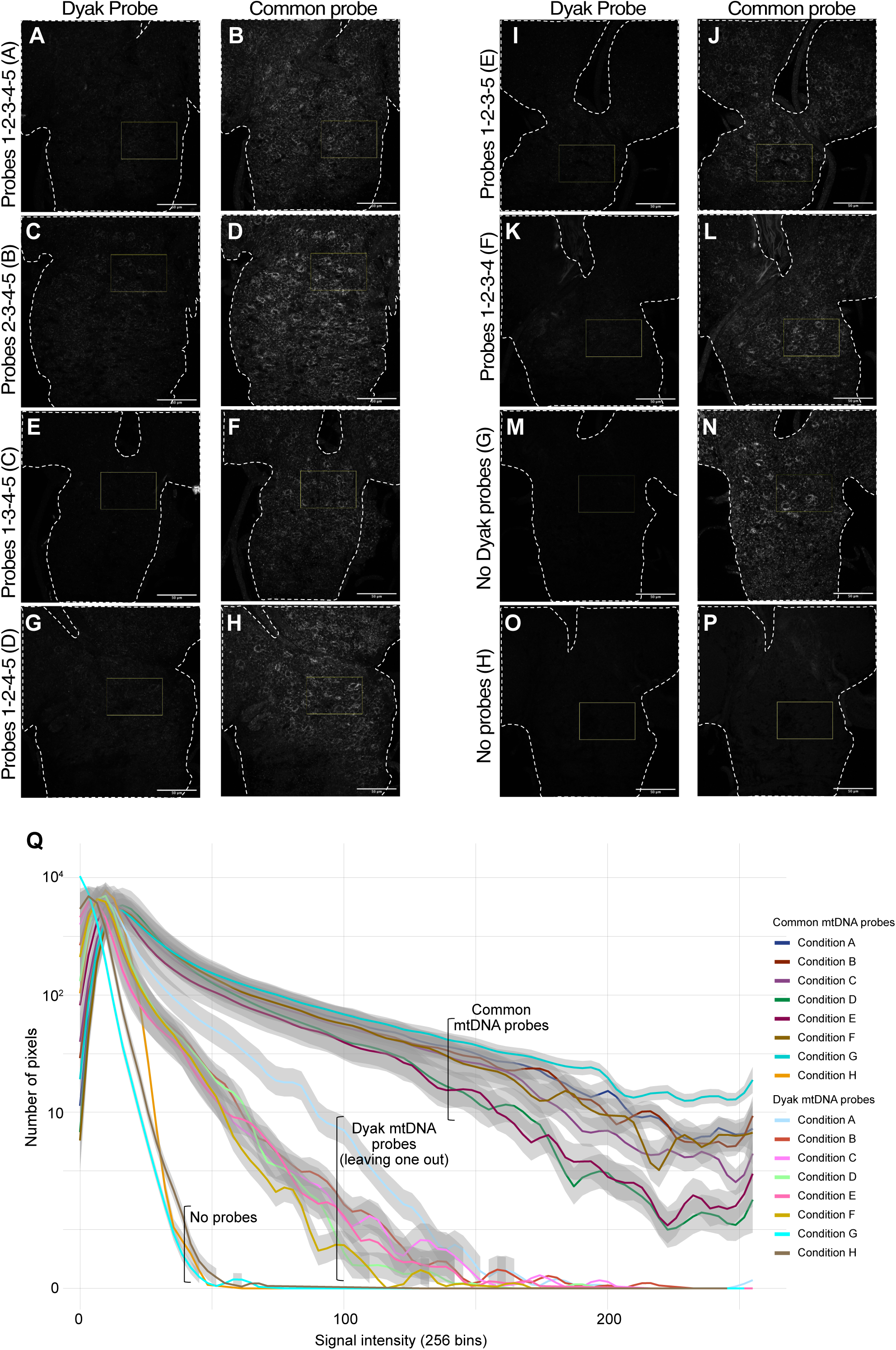
Specificity of individual probes for *D. yakuba* mtDNA-smFISH. To assess background of the probe-set specific to *D. yakuba* mtDNA, each of the five individual probe pairs was removed separately from the probe mix used for mtDNA-smFISH on brains from *Drosophila* with homoplasmic *D. melanogaster* mtDNA. Removing individual *D. yakuba* probes (representative images in A,C,E,G,I,K, light colours in Q), all resulted in comparable background levels. mtDNA-smFISH with the common probe set was used as a positive control (representative images in B,D,F,H,J,L, dark colours in Q). No background or signal was observed in absence of *D. yakuba* probes (M,Q) or any probe (O,P,Q). Scale bars are 50μm.

**Supplementary figure 8:**
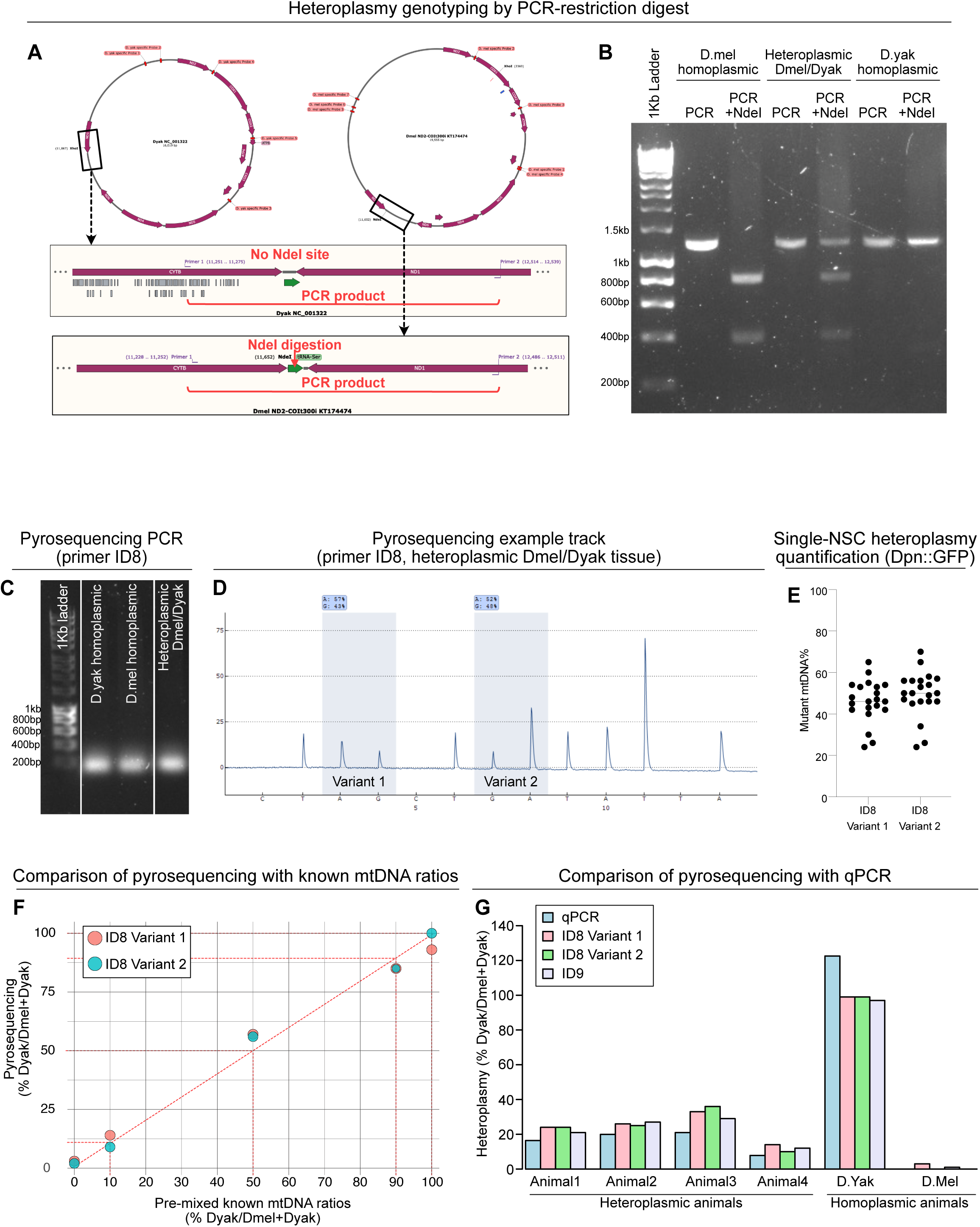
Heteroplasmy measurement with differential restriction digestion or pyrosequencing. (A,B) Schematic (A) and representative gel image (B) of PCR amplification and NdeI digestion to distinguish *D. yakuba* (WT) from *D. melanogaster* (mutant) mtDNA, with *D. yakuba* lacking the NdeI site. PCR amplification produced a 1284bp (*D. melanogaster*) or 1289bp (*D. yakuba*) product, which, upon digestion with NdeI, resulted in two bands of 425bp and 859bp for the *D. melanogaster* genome only. (C-G) Optimisation of pyrosequencing to distinguish *D. yakuba* (WT) from *D. melanogaster* (mutant) mtDNA, using primer pairs ID8 (C-G) or ID9 (G). PCR using primer pair ID8 amplifies mtDNA from homoplasmic or heteroplasmic *Drosophila* strains with comparable efficiency (C), and allows quantification of heteroplasmy levels using two independent variants in the same PCR fragment (D). Pyrosequencing detects heteroplasmy levels in single NSCs sorted from Dpn::GFP third instar larvae (E). Pyrosequencing detects known ratios of mixed mtDNA from homoplasmic *D. melanogaster* and *D. yakuba* (F), and detects heteroplasmy levels from heteroplasmic or homoplasmic *Drosophila* with comparable accuracy to a previously validated qPCR method ^61^ (G).

**Supplementary Figure 9:**
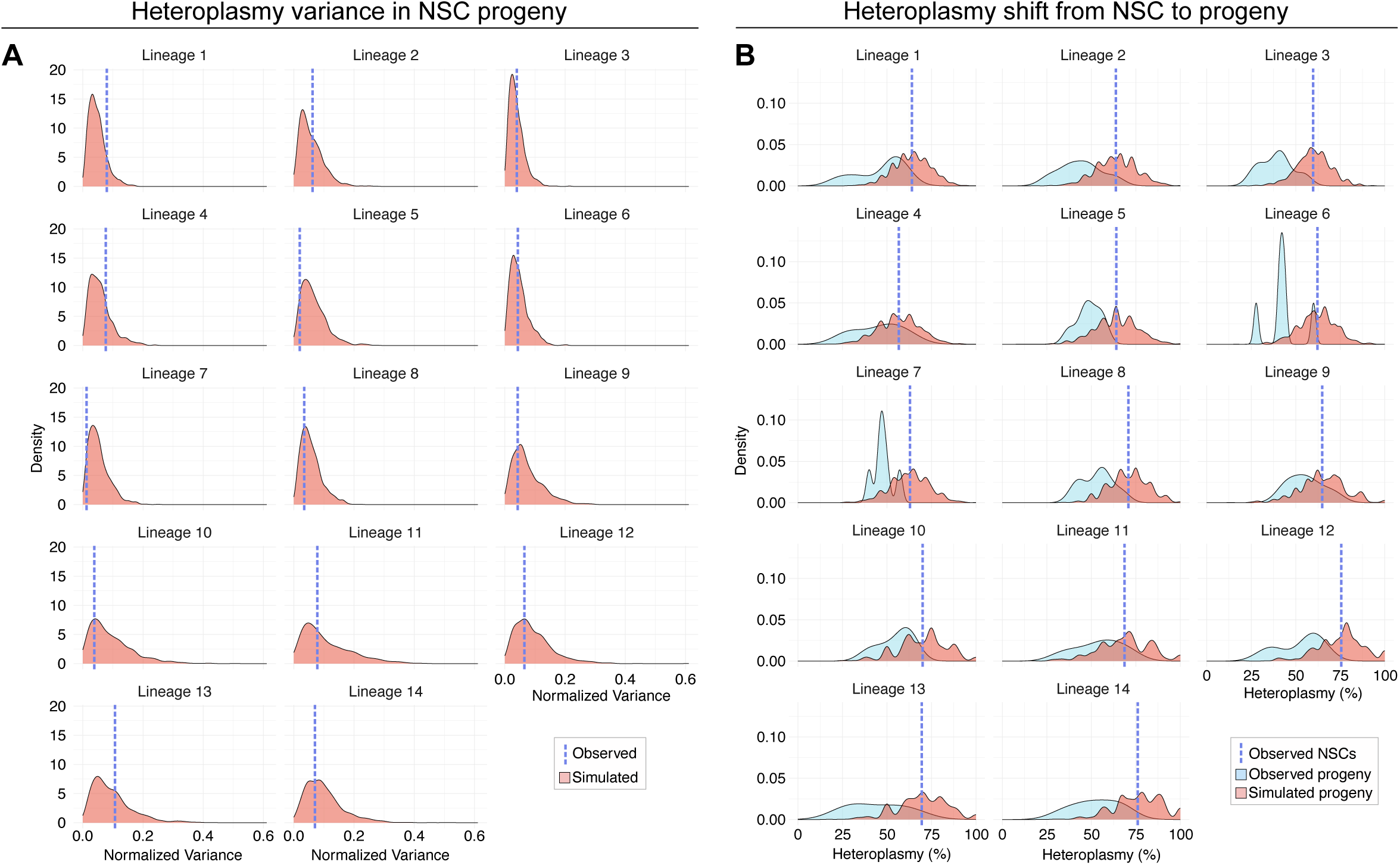
Modelling of mtDNA segregation through a somatic bottleneck. (A) Normalized heteroplasmy variance in progeny cells per lineage (red) with the dashed line representing the normalized heteroplasmy variance in observed progeny cells. (B) Heteroplasmy levels for observed progeny cells (light blue) and simulated progeny cells (red) per lineage. The dashed line (dark blue) represents the heteroplasmy level in the parent NSC in each lineage.

**Supplementary Table 1: smFISH-mtDNA probe sequences.**

